# *OmicsNavigator*: An auditable scientific partner for scalable hypothesis validation in spatial omics

**DOI:** 10.1101/2025.07.21.665821

**Authors:** Yiyao Li, Nirvi Vakharia, Weixin Liang, Aaron T. Mayer, Ruibang Luo, Alexandro E. Trevino, Zhenqin Wu

## Abstract

Translating high-dimensional, spatially resolved molecular datasets into testable biological findings remains a major research bottleneck. Here, we present *Omic-sNavigator*, an autonomous large language model-powered system for end-to-end data exploration and hypothesis validation on spatial omics data. *OmicsNaviga-tor* reasons directly over the multi-modal inputs of spatial omics data, including visual and molecular signatures, to perform knowledge-guided annotation of spatial structures. We show that by transforming high-dimensional data into textual interpretations, *OmicsNavigator* enables zero-shot semantic retrieval of tissue biomarkers and the reconstruction of patient-level disease profiles from raw omics observations. Furthermore, *OmicsNavigator* features an objective hypothesis validation engine governed by pre-registered, human-audited blueprints. By validating the system across datasets spanning diverse pathological conditions including diabetic kidney disease, kidney transplant rejection, and COVID-19 pulmonary pathology, we demonstrate that *OmicsNavigator* generates evidence-based, human-readable insights from spatial omics data, with potential to accelerate spatial biology discoveries.

## 1 Introduction

Spatial omics technologies characterize the detailed molecular composition of tissues within their native architecture, advancing our understanding of complex tissue biology [1]. Platforms such as 10x Visium [2], MERFISH [3], Xenium [4], CODEX/PhenoCycler [5], and MIBI-TOF [6] now routinely generate terabytes of data comprising spatially resolved, high-dimensional molecular measurements for diverse tissue landscapes. However, this technological leap has precipitated an analytical bottleneck: while we possess maps of these territories, we lack the autonomous navigators to traverse them.

Significant progress has been made in the computational modeling of spatial omics data. Current state-of-the-art approaches extract cellular features—such as gene/protein expression and morphology—represented as matrices [7–10] or graphs [9, 11–13]. These representations feed into advanced artificial intelligence models to generate latent embeddings for diverse downstream analyses. Concretely, specialized tools have now robustly addressed key analytical tasks such as cell segmentation [14, 15], cell type clustering [16–18], and neighborhood annotation [19, 20], enabling the curation of detailed cellular maps of tissue architectures. Nevertheless, an open challenge remains: how to seamlessly bridge these computational abstractions with broader clinical contexts and pre-existing biological knowledge. Consequently, the field faces an “interpretation gap,” where the richness of spatial data is often reduced to catalogs of cells and neighborhoods. While powerful, these approaches risk devaluing intuitive, human-readable insights needed to inform biological understanding and discoveries.

The integration of Large Language Models (LLMs) into bioinformatics has offered promising solutions for automated pipeline orchestration, code generation, and tool execution [21–27]. These systems demonstrated strong capabilities in executing existing cell-centric pipelines—composing and executing code, and interfacing with existing bioinformatics tools. An area of keen interest that remains underexplored is enabling agentic systems to operate and reason directly over the underlying multi-modal data— such as interpreting local molecular signatures alongside tissue morphology to identify pathological structures associated with patient phenotypes. Expanding agentic capabilities into this space is essential to fully leverage the rich spatial contexts—visuals of tissue morphology, subcellular organization, and surrounding environment of cells— that are crucial for biological interpretation. To unlock the full potential of spatial omics and bridge the interpretation gap, we envision a next-generation spatial biology computational tool that moves beyond the cell-centric task space and toward the autonomous extraction and synthesis of evidence, grounded in the spatially resolved data, the biological prior knowledge, and the study-specific research questions.

In parallel, the rapidly increasing capabilities of AI agents in scientific research raise concerns about the reliability of their output. State-of-the-art agentic scientific tools have enabled the automated, parallel generation of thousands of ideas, hypotheses, and candidate insights [28–31]. While this throughput is a testament to the power of LLM-driven discovery, it simultaneously overwhelms researchers with findings that may stem from spurious correlations or statistical artifacts [31, 32]. Existing pipelines, whether led by human researchers or AI agents, remain susceptible to confirmation bias and subjective parameter tuning—issues that are particularly relevant in spatial biology, where studies typically involve small biological cohorts with thousands of profiled cells per sample, and where subjective selection of neighborhoods might alter analysis conclusions. We argue that to serve as a genuine research partner in this high-throughput era, a next-generation computational tool must follow a framework with autonomous self-constraints, complemented by human oversight.

To realize this vision of a rigorous, spatially aware scientific partner for spatial biology, we introduce *OmicsNavigator* (Figure 1), an autonomous agentic system designed for objective, end-to-end data exploration and hypothesis validation. *OmicsNavigator* demonstrates the following key capabilities: (1) high-throughput, multi-modal interpretation of spatial structures by directly reasoning over visual morphology, molecular signatures, and curated biological prior knowledge; (2) zero-shot retrieval and discovery of spatial biomarkers using natural language queries; (3) study-scale summary and reasoning to derive disease-relevant insights; and most crucially, (4) autonomous, self-constrained hypothesis validation, enforced by human-in-the-loop (HITL) audits and pre-registered evaluations. To achieve these capabilities in practice, we developed and integrated novel mechanisms, including a specialized Plan Module that synthesizes literature and user queries into Background Knowledge Checklists and Verification Blueprint, a dual-channel profiling mechanism for annotations of spatial structures, a “Cluster-Anchor-Expand” mechanism for efficient retrieval, and a directed acyclic graph (DAG)-guided hypothesis validation pipeline.

**Fig. 1.**
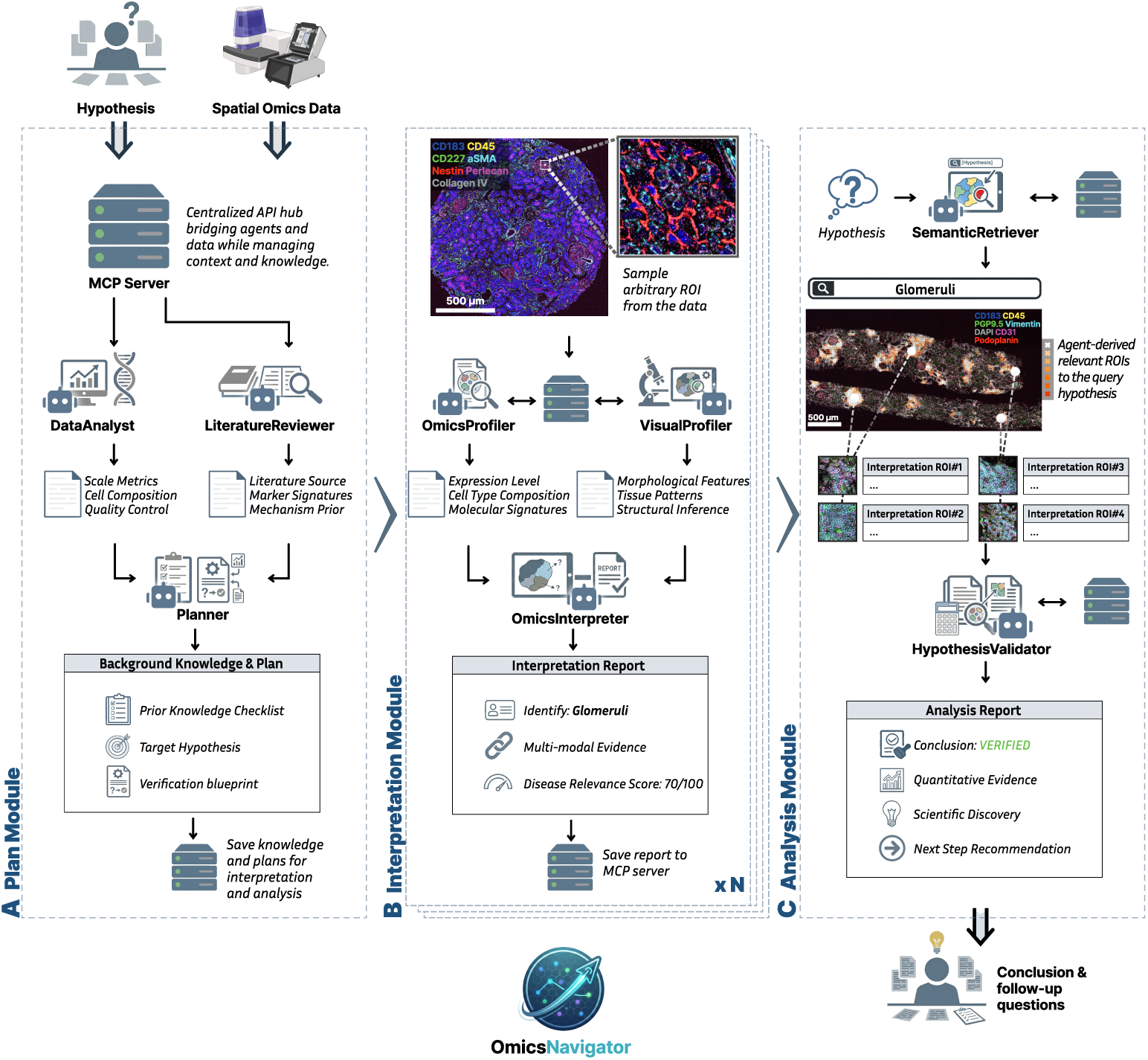
*OmicsNavigator* architecture: A modular autonomous agentic system for end-to-end spatial omics data exploration and hypothesis validation. (a) The Plan Module. A central *Planner* Agent parses natural language queries and hypotheses. It establishes domain-specific biological priors by coordinating a *LiteratureReviewer* for knowledge extraction and a *DataAnalyst*for initial profiling of data baselines. (b) The Interpretation Module. This module introduces a dual-channel profiling mechanism to prompt a *VisualProfiler* that extracts visual morphology features and an *OmicsProfiler* that extracts molecular signatures as structured natural language descriptions. These streams are subsequently synthesized by the *OmicsInterpreter*, which utilizes domain-specific biological priors to provide interpretations. (c) The Analysis Module. This module contains a *SemanticRetriever* to conduct retrieval of hypothesis-related spatial structures, and a *HypothesisValidator* to synthesize preceding outputs and perform statistical validations. Following the execution plan (i.e., Verification Blueprint) established by the *Planner*, this module generates a detailed report to validate or refute user hypothesis.

Across diverse datasets spanning diabetic kidney disease (DKD) [33], kidney transplant rejection [34, 35], and COVID-19 pulmonary pathology [36], *OmicsNavigator* consistently demonstrates reliable performance in data interpretation, exploration, and retrieval. It delivers evidence-backed verifications and refutations for user-specified hypotheses, and demonstrates capabilities to discover new biological narratives beyond initial queries. By enabling researchers to interact directly with high-dimensional tissue landscapes through natural language, *OmicsNavigator* transforms technically demanding computational workflows into an intuitive, human-readable process.

## 2 Results

### 2.1 Autonomous Curation of Biological Priors

Interpretations of spatial omics data, an essential step towards automated scientific discovery, rely heavily on domain-specific biological prior knowledge. Acquiring such priors, however, often demands manual curation by domain experts, creating a bottleneck in translating computational analysis to biological understanding. As LLMs are increasingly integrated into modern computational biology workflows for omics analysis[37–40], this need for biological priors becomes even more critical, as LLMs without strict evidence-based guardrails are prone to hallucinations and misalignment. To overcome this expert-dependency bottleneck and ground LLM reasoning in preexisting biological knowledge, *OmicsNavigator* introduces a dedicated Plan Module (Figure 1A) to autonomously curate these domain-specific biological priors before any close examination begins.

This autonomous knowledge curation is executed via a sequential workflow of specialized sub-agents to extract and synthesize evidence from diverse sources. First, a *DataAnalyst* sub-agent performs a global, unsupervised profiling of the input spatial omics data. It establishes baseline metrics such as cell composition and spatially variable expression. A *LiteratureReviewer* is then deployed to extract existing evidence from the literature. It translates the data baselines and user inputs into structured semantic queries, and then executes targeted queries across publication repositories (e.g., PubMed) and biological databases (e.g., NCBI [41], QuickGO [42]), focusing on mining relevant pathological mechanisms and known biomarkers.

Based on these priors—background knowledge, relevant literature, and dataset baselines—a *Planner* agent synthesizes them into structured Background Knowledge Checklists (Appendix B.2). These checklists encode the reasoning logic of a human expert, providing domain-specific criteria for subsequent analysis. For instance, in the context of DKD, checklists include morphological guidelines (e.g., identifying a glomerulus via a distinct Perlecan/Collagen ring), molecular enrichment rules (e.g., differentiating tubular structures from glomeruli by epithelial protein markers), disease-associated phenotypes (e.g., fibrosis and inflammation), and quantitative indicators for disease severity. By establishing these priors, the checklists serve as a cognitive foundation that constrains the downstream interpretations and reasoning, thereby reducing hallucinations.

Beyond establishing biological priors, the Plan Module simultaneously lays the groundwork for hypothesis validation. In addition to the checklists, the central *Planner* agent parses the user-supplied queries and proposes a standardized Verification Blueprint (Appendix B.3). The blueprint defines exact hypothesis-testing pipelines before any analysis is run—locking in target variables, inclusion criteria, and testing paths in the format of a DAG (Supplementary Figure 1).

Crucially, before executing subsequent analysis, *OmicsNavigator* requires a user review of the checklists and the blueprint. In this step, researchers can revise and augment these documents; once confirmed, they are locked throughout the analysis to establish a fixed execution space and to serve as a guardrail against agentic confirmation bias.

### 2.2 Intelligent Multi-modal Interpretation of Spatial Structures

Data exploration, hypothesis validation, and insight discovery in spatial omics studies all build on accurate interpretations of spatial structures across scales—cells, neighborhoods, and beyond. To achieve that, *OmicsNavigator* employs a dedicated interpretation module that leverages the established domain-specific biological priors to interpret raw observations.

To analyze structures of diverse scales, *OmicsNavigator* interacts with the complex, high-dimensional spatial omics data using units termed Regions of Interest (ROIs). Operationally, an ROI is defined by a local bounding box of variable size centered on a specific spatial coordinate or cell, capturing both microscopic morphology and molecular profiles within that window. For any given ROI, the interpretation module employs a dual-channel profiling mechanism to characterize it: two specialized sub-agents— a *VisualProfiler* for visual morphology images and an *OmicsProfiler* for molecular profiles—extract features as natural language descriptions. Both sub-agents are guided by the pre-registered Background Knowledge Checklists. The two streams of information are subsequently fused by the *OmicsInterpreter* (Figure 2A), which integrates the evidence with the biological priors to produce a spatially aware interpretation of the structures contained in the queried ROI.

**Fig. 2.**
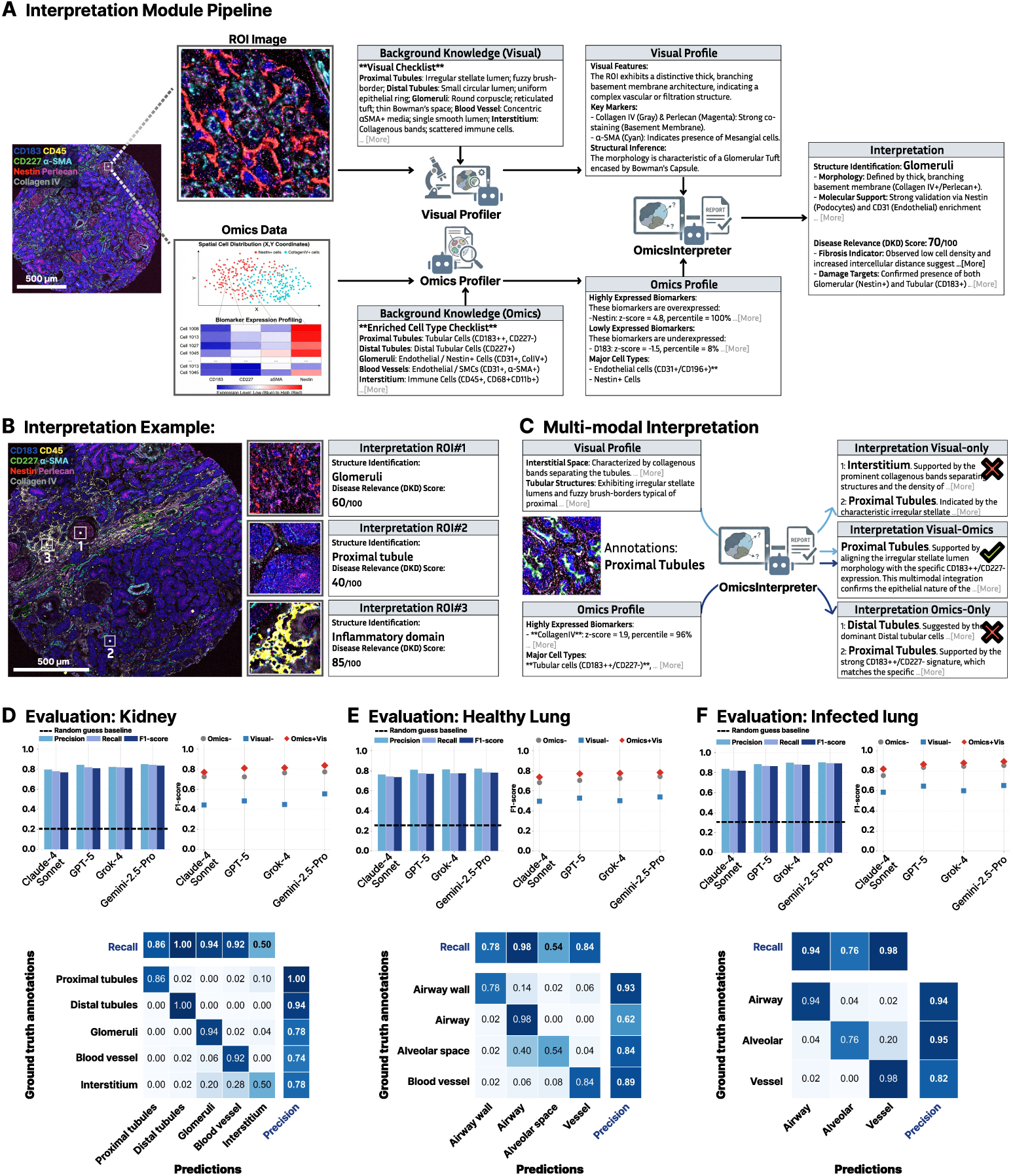
Intelligent multi-modal interpretation via dual-channel profiling and fusion. (a) Schematic of the interpretation module’s pipeline. The dual-channel profiling mechanism extracts descriptive features from two parallel input streams—visual morphology and molecular profiles—which are subsequently fused. This schematic also highlights the injection of Background Knowledge Checklists to ground agentic reasoning. (b) Representative annotations of a human kidney tissue section. *OmicsInterpreter* accurately identifies complex structures such as glomeruli, tubular structures, and disease-induced inflammation domains. (c) Ablation study demonstrating the contribution of multi-modality. Uni-modal baselines exhibit specific failure modes: visual-only interpretation (top) confuses early interstitial fibrosis with edema due to morphological similarity; Omics-only interpretation (bottom) is confused by the overlapping biomarker expression. The fused interpretation by *OmicsInterpreter* (center) correctly synthesizes both cues to derive the correct conclusion. (d-f) Quantitative benchmarking. F1 scores for zero-shot structural annotations against human labeled ground truth on diabetic kidney disease (DKD) (d), healthy lung (e), and SARS-CoV-2-infected lung (f) datasets.

The prior-guided design allows *OmicsNavigator* to produce interpretations with clear biological relevance, including explicit linkage to the disease pathology. As demonstrated in the annotations of a late-stage DKD kidney tissue section (Figure 2B), *OmicsInterpreter* successfully performed interpretations for three arbitrarily sampled ROIs across the heterogeneous tissue landscape. It correctly identified canonical structures (e.g., glomeruli) and disease-induced inflammation domains. Furthermore, it assigned high relevance scores to the glomeruli and inflammation ROIs, consistent with their pathological significance in DKD.

Integration of multi-modal information is another core feature of the interpretation module. An illustrative example is shown in Figure 2C. Visual-only interpretation of this ROI leaned towards interstitium due to morphological clues and irregular luminal shapes. Conversely, omics-only interpretation misclassified the ROI as distal tubules due to protein marker profiles. Such failures were often observed in ambiguous regions, where the fused interpretation generated by *OmicsInterpreter* resolves the ambiguity and correctly annotates the queried ROI.

To quantify interpretation performance, we evaluated zero-shot annotation accuracy against ground truth human annotations across three datasets of diverse biological contexts—DKD-affected kidney, healthy lung, and SARS-CoV-2-infected lung (Figure 2D–F). Across all scenarios, *OmicsNavigator*’s interpretations consistently achieved strong F1 scores in annotating canonical structures. Furthermore, the multi-modal configuration consistently outperformed both visual-only and omics-only baselines. The incorporation of biological priors proved critical for accurate interpretations (Supplementary Figure 2). Minor differences were observed between agents powered by different LLM engines, among which Gemini-2.5-Pro yielded the highest overall interpretation performance. Confusion matrices confirmed good precision in distinguishing canonical structures (e.g., proximal vs. distal tubules, airway walls vs. alveolar spaces). We adopt this Gemini-driven configuration as the standard setup for all subsequent experiments.

We emphasize that the ROI-level interpretations performed by *OmicsNavigator* are different from the related task of spatial domain/niche discovery, which focuses on unsupervised clustering of observations. Here, interpretations are generated independently for each ROI and are grounded in biological prior knowledge. This enables more flexible and nuanced characterization of rare or disease-associated structures, and yields semantically meaningful annotations that directly support subsequent reasoning about patient-level disease states.

### 2.3 Scalable Zero-shot Semantic Retrieval of Spatial Biomarkers

Building on accurate interpretations of ROIs, we next introduce a *SemanticRetriever* agent that performs semantic retrieval over spatial omics studies. Concretely, given a user query or hypothesis, *SemanticRetriever* identifies and locates all ROIs containing query-relevant structures or spatial biomarkers and characterizes their properties in relation to the query.

A naive solution would apply the interpretation module exhaustively to every ROI in the study, followed by searching the user query against the resulting interpretation text corpus. This is infeasible in practice due to computational and budgetary constraints, as spatial omics studies routinely contain millions of cells. To enable accurate retrieval at scale, *SemanticRetriever* leverages a “Cluster-Anchor-Expand” pruning strategy for efficient, zero-shot spatial biomarker discovery.

The pipeline proceeds in three stages (Figure 3 A). First, ROIs are enumerated and grouped based on their multi-modal features and spatial coordinates, using a multi-level clustering strategy inspired by previous works [43, 44]. We then prune each resulting group using a distance-weighted stochastic sampling approach, resulting in biologically representative “Anchor ROIs”. Typically for each tissue section, this approach yields around 50-100 anchor ROIs, which are then submitted to the interpretation module. Subsequently, the user query, parsed as natural language keywords, is compared against anchor ROI interpretations using text-based similarity metrics to retrieve top matches, termed “hits”. Lastly, *SemanticRetriever* employs an expansion step that leverages hits to identify all relevant ROIs across the study. More details of this pipeline can be found in Methods.

**Fig. 3.**
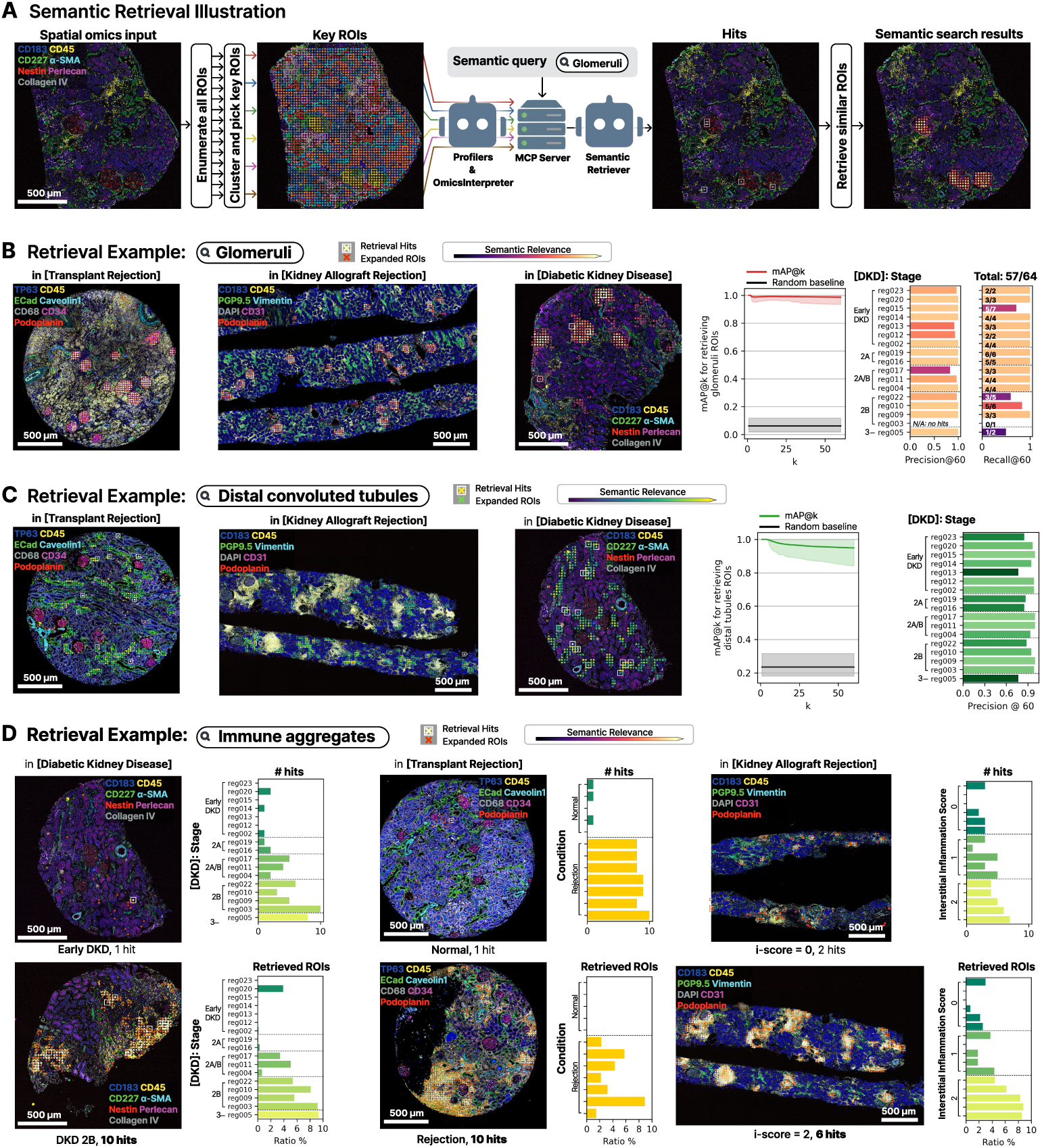
High-throughput semantic retrieval via the Cluster-Anchor-Expand strategy. (a) Schematic of the retrieval engine. To process large-scale data, the system groups ROIs, interprets representative ROIs, retrieves hits relevant to user queries, and uses them as anchors to expand semantic tags to cover the entire study. (b-c) Retrieval examples for canonical structures of glomeruli (b) and distal convoluted tubules (c) across three different spatial omics studies from different institutions, pathologies, and biomarker panels. Mean average precision at k (mAP@k) curves, precision@60, recall@60 metrics are calculated on the DKD cohort, where *OmicsNavigator* achieved strong performances. (d) Discovery of disease-associated spatial biomarkers using targeted queries for “Immune Aggregates”. The quantities of retrieved ROIs correlate strongly with pathologist-assigned stages or inflammation levels across all three studies.

We evaluated *SemanticRetriever* using two canonical structures as spatial biomarkers: glomeruli and distal convoluted tubules. Here, the target structures are queried in plain natural language against three kidney spatial biology datasets spanning different institutions, pathologies, and biomarker panels. Qualitatively, the retrieved ROIs across datasets demonstrated consistent and close alignment with expert annotations (Figure 3). In the DKD dataset, *SemanticRetriever* achieved a precision of 0.97 at the top-60 (*k* = 60) cutoff when retrieving glomeruli, with 89% of all glomeruli in the study identified at this cutoff (Figure 3 B). Queries for distal convoluted tubules yielded a precision of 0.93 at the same *k* = 60 cutoff (Figure 3 C). The mean Average Precision (*mAP* @*k*) curves for both cases remained near 1.0 with tight 90% confidence intervals, indicating both high accuracy and stability of our approach.

In addition to canonical structures, *SemanticRetriever* can also discover disease-associated spatial biomarkers using natural language queries. When queried for “Immune Aggregates”, a hallmark of various pathological conditions, *SemanticRetriever* located ROIs that exhibit strong enrichment in major immune cell stains (e.g., CD45, CD68). Furthermore, the abundance of retrieved ROIs demonstrated strong positive correlations with clinically determined staging or inflammation scores assigned by pathologists (Figure 3 D). Together, these results establish that the system can locate pathologically meaningful spatial structures in the study based on natural language descriptions alone, enabling further quantitative analyses across regions and patients.

### 2.4 Patient-Scale Disease State Characterization

Patient-level pathological phenotypes typically emerge from the collective accumulation of micro-scale aberrations distributed across the tissue. Such micro-scale pathological changes can be captured and located using *OmicsNavigator*’s ROI interpretation and retrieval capabilities demonstrated above. To further enable patient-level clinical analysis, it is necessary to aggregate these local observations into coherent disease state summaries. To this end, *OmicsNavigator* implements an aggregation pipeline that reconstructs patient-level disease states from ROI-level observations and interpretations.

The aggregation pipeline proceeds in three major steps. First, *OmicsNavigator* identifies and retrieves disease-relevant ROIs. This can be performed by targeting known pathological structures via *SemanticRetriever*, as demonstrated in Figure 3 D. Alternatively, thresholding on the disease relevance scores assigned by the interpretation module (e.g., score ≥ 80) can surface all ROIs that are relevant for evaluating the patient’s disease state. Second, *OmicsNavigator* counts these disease-relevant ROIs to derive quantitative measures of how much the patient sample is characterized by disease-induced changes. Third, *OmicsNavigator* synthesizes the natural language interpretations of these disease-relevant ROIs to compose a patient-level summary that captures the core pathological changes at play.

We applied this aggregation pipeline to the DKD cohort to reconstruct clinical progression (Figure 4A). Counts of highly disease-relevant ROIs remained near zero for early-stage samples but increased substantially in late-stage samples, yielding a statistically significant, stepwise increase across stages (ANOVA followed by Tukey’s HSD, *p <* 0.001). Concurrently, the synthesized semantic summaries reflected shifts from localized structural changes with minimal inflammation to severe immune infiltration combined with interstitial fibrosis and vascular rarefaction. In another kidney cohort of transplant rejection (Figure 4B), counts of disease-relevant ROIs differed significantly between acute rejection samples and normal controls (Student’s t-test, *p <* 0.001). Crucially, the normal baseline yielded zero hits, highlighting the system’s high specificity. The corresponding semantic summaries described the immune quiescence and structural integrity in normal samples, contrasted with the severe host-versus-graft damage in acute rejection samples.

**Fig. 4.**
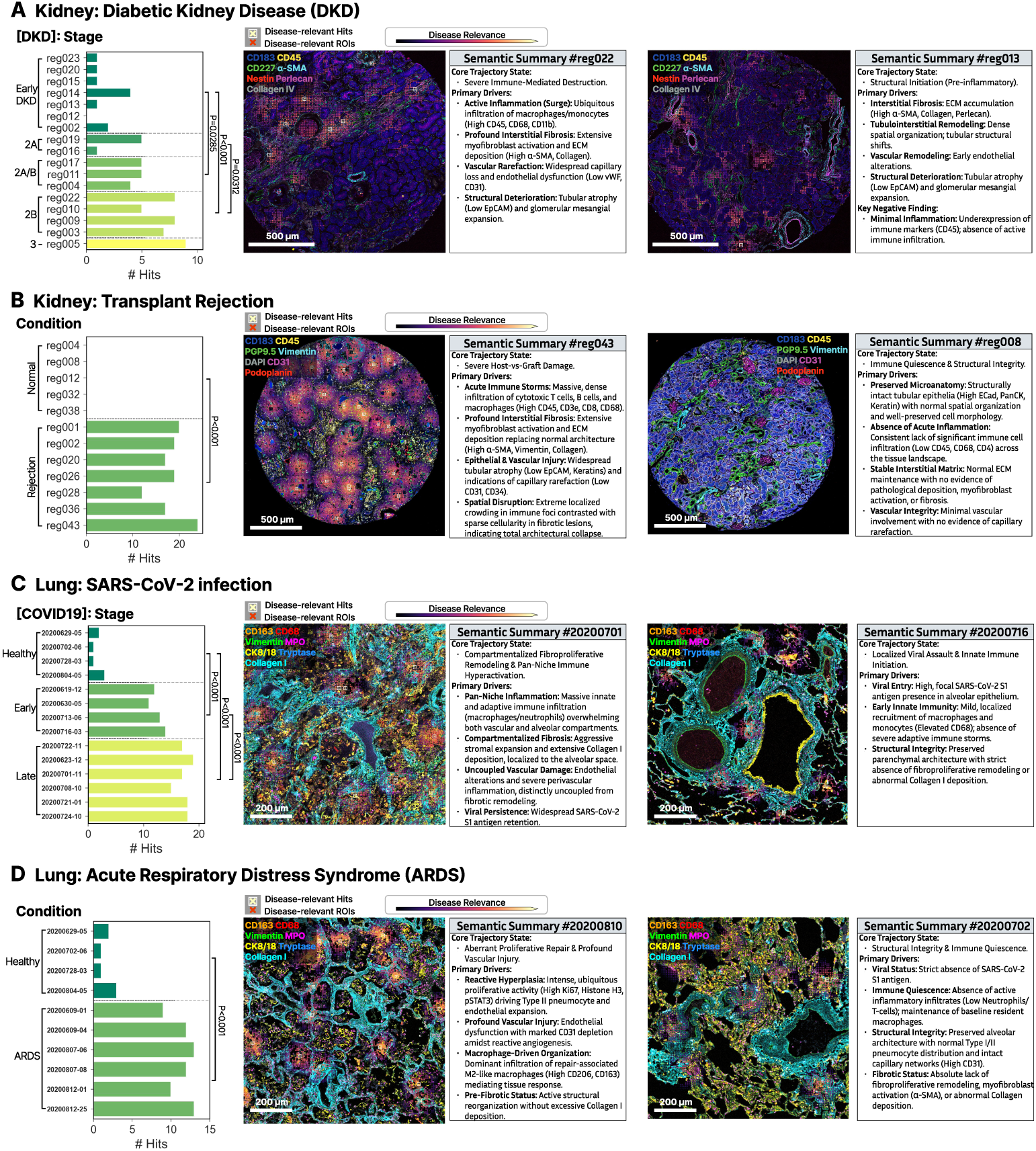
Patient-level disease state reconstruction and clinical stratification. (a) Progression of DKD across regions. By executing an aggregation pipeline on disease-relevant ROIs (relevance score *≥* 80), *OmicsNavigator* tracks the collective accumulation of disease severity. The quantitative count shows a statistically significant, stepwise increase from early-stage to late-stage (*p <* 0.001). Synthesized patient-level summaries capture the biological transition from initiation of structural degradation to severe immune-mediated destruction along with interstitial fibrosis. (b) Identification of kidney changes due to transplant rejection. *OmicsNavigator* distinguishes regions from acute rejection patients from normal baselines (*p <* 0.001). The semantic summaries explicitly contrast the immune quiescence and structural integrity in healthy tissues against severe host-versus-graft damage dominated by acute immune storms in transplant rejection samples. (c) Characterization of pulmonary SARS-CoV-2 infection. *OmicsNavigator* maps the progressive accumulation of disease-relevant ROIs, establishing a severity gradient from the healthy baseline to early and late-stage infection (*p <* 0.001). Semantic summaries identify the shift from localized early viral assault to pan-tissue macrophage enrichment and compartmentalized alveolar fibrosis. (d) Identification of pathological changes in acute respiratory distress syndrome (ARDS). *OmicsNavigator* separates ARDS samples from healthy base-lines (*p <* 0.001). Its semantic summary identifie2s5aberrant proliferative repair, vascular injury, and macrophage enrichment. Statistical analysis: For multi-group comparisons (panels a and c), statistical significance is determined using a one-way ANOVA followed by Tukey’s HSD post-hoc test. For two-group comparisons (panels b and d), significance is assessed using an independent Student’s t-test.

We next applied *OmicsNavigator* to two lung disease conditions (Figure 4C, D). In the COVID-19 cohort, the system captured a significant severity gradient from the healthy baseline to early infection to late-stage exacerbation (ANOVA followed by Tukey’s HSD, *p <* 0.001). Semantic summaries identified a pathological progression from localized early viral injury to widespread pathogenesis: pan-tissue immune activation and compartmentalized alveolar fibrosis. In the comparison between acute respiratory distress syndrome (ARDS) and healthy samples, *OmicsNavigator* differentiated the two phenotypes (Student’s t-test, *p <* 0.001). The semantic summary identified aberrant proliferative repair, vascular injury, and macrophage-driven organization as core drivers of the ARDS organizing phase.

These reconstructed disease trajectories are deterministic by design. The criteria for targeting ROIs—biological priors of the disease and relevance scoring rubrics—are pre-registered in the Background Knowledge Checklists before any analysis is run. Consequently, this approach mitigates potential confirmation bias during patient-level reasoning.

### 2.5 Pre-registered Autonomous Hypothesis Validation

Finally, to translate high-dimensional spatial omics data into scientific deliverables, *OmicsNavigator* leverages a *HypothesisValidator* agent that integrates outputs from all components to perform rigorous, end-to-end hypothesis validation (Figure 5). *HypothesisValidator* inherits outputs from preceding phases: (1) the Verification Blueprint from the *Planner* agent (Section 2.1) serves as an execution guide; (2) zero-shot semantic retrieval (Section 2.3) of hypothesis-related ROIs (e.g., glomeruli for kidney function studies, alveoli for pulmonary state analysis) by *SemanticRetriever* provides pools of relevant ROIs for targeted analysis; (3) interpretations (Section 2.2) by *OmicsInterpreter* offer detailed characterization of these ROIs; and (4) patient-level aggregation (Section 2.4) synthesizes the ROI-level data into patient-level observations across different pathological conditions for statistical assessment.

**Fig. 5.**
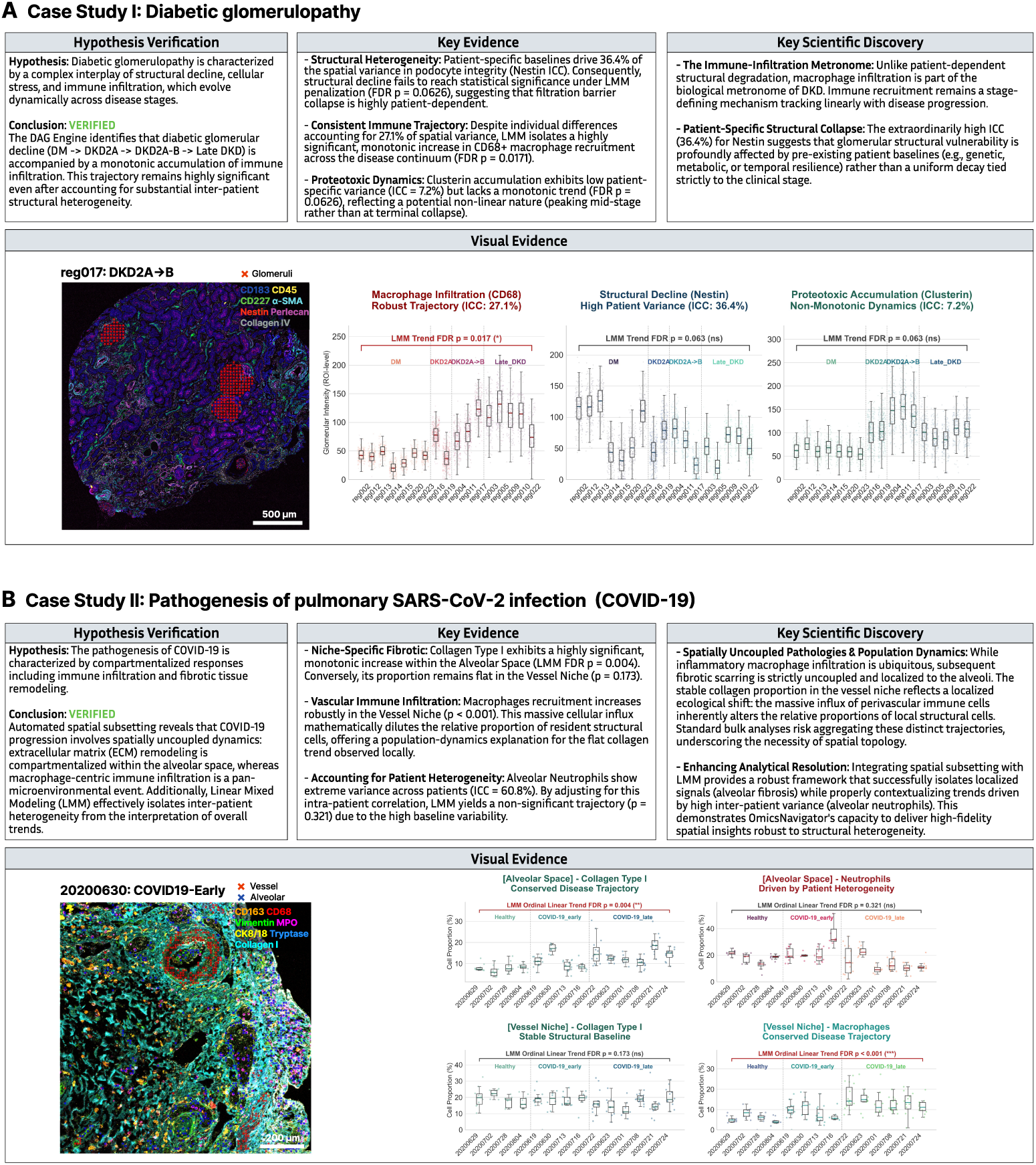
Autonomous hypothesis validations of spatially resolved disease dynamics. (a) Hypothesis validation regarding diabetic glomerulopathy in the DKD study. The system processed a natural language hypothesis regarding the monotonic trend of structural decline, cellular stress, and immune infiltration. Operating under the Verification Blueprint, *HypothesisValidator* utilized a Linear Mixed Model (LMM) to model the stage-dependent dynamics. It provided data-driven insights, identifying CD68+ macrophage recruitment as a highly significant, monotonic trajectory, while suggesting that structural decline (Nestin) was heavily influenced by inter-patient variance. Trend plots mapping these markers across disease stages were automatically generated to support the findings. (b) Hypothesis validation regarding compartmentalized pathogenesis in the COVID-19 pulmonary tissue study. The system executed retrieval to decouple alveolar space and vascular ROIs. *HypothesisValidator* found that fibrotic scarring is compartmentalized to the alveolar space, while macrophage enrichment is a ubiquitous, pan-compartmental event. Alveolar neutrophils showed high heterogeneity (ICC = 60.8%); after adjustment, the patient-level association became non-significant. Plots were automatically generated to support the findings.

Here, we demonstrate *HypothesisValidator*’s capability using two case studies across different organ systems and datasets. In both cases, *HypothesisValidator* follows the pre-registered DAG defined in the Verification Blueprint to execute the computational pipeline. The frameworks proposed autonomously by the *Planner* agent utilize Linear Mixed Models (LMMs) to account for heterogeneity of ROIs nested within individual patients and apply multiple-testing corrections (e.g., Benjamini-Hochberg False Discovery Rate (FDR)) to control false positives.

#### Case Study I: Diabetic Glomerulopathy

We first evaluated whether three aspects of diabetic glomerulopathy—structural decline, cellular stress, and immune infiltration—progress monotonically with DKD stage (Figure 5A). The *Verification Blueprint* formulated an analytical protocol targeting glomeruli, and utilized the LMM engine to account for patient-level variances. Guided by the DAG, the *HypothesisValidator* quantified the trajectories of key spatial biomarkers and protein expression relevant to glomerulopathy across stages.

*HypothesisValidator* produced detailed, data-driven findings that describe the changes in glomeruli during DKD progression: it found that structural decline, measured by the podocyte integrity marker Nestin, is highly patient-dependent. Patient-specific baselines accounted for 36.4% of the spatial variance (Nestin intraclass correlation (ICC)). Consequently, the overall trend of structural decline did not reach statistical significance under LMM adjustment (FDR *p* = 0.063) and was therefore refuted. This result suggests that the collapse of the filtration barrier is heavily influenced by patient-specific baseline factors rather than being uniformly tied to clinical stage. A larger, better-powered study is needed to disentangle these patient-level effects from stage-dependent progression.

In contrast, the LMM detected a significant, monotonic increase in CD68+ macrophage recruitment across the disease continuum (FDR *p* = 0.017) despite moderate individual differences (ICC = 27.1%). This supports immune infiltration as a conserved mechanism that increases monotonically with DKD progression, regardless of a patient’s glomerular structural baseline. Finally, proteotoxic stress, measured by glomerular Clusterin levels, exhibited low patient-specific variance (ICC = 7.2%) and a non-monotonic trend. While the hypothesis of a monotonic trend was later refuted (FDR *p* = 0.063), *HypothesisValidator* noted a potential non-linear dynamic that peaks at mid-stage as a candidate for further investigation.

*OmicsNavigator*’s re-analysis converges with Kondo et al. [33] on the major findings: a monotonic increase in CD68+ macrophage infiltration across DKD progression, and pronounced heterogeneity in glomerular structural decline. The LMM engine employed by *HypothesisValidator* quantifies this heterogeneity explicitly—Nestin’s high patient-level ICC (36.4%) provides support for the patchiness that the original study described visually. Additionally, we characterize a potential non-monotonic trajectory of cellular stress that peaks at mid-stage DKD, a dynamic not reported in the original analysis.

#### Case Study II: Pathogenesis of Pulmonary SARS-CoV-2 Infection (COVID-19)

We next applied *OmicsNavigator* to investigate pulmonary pathogenesis during COVID-19, testing whether the disease is characterized by spatially compartmentalized responses including fibrosis and inflammation (Figure 5B). Guided by the *Verification Blueprint*, *HypothesisValidator* executed automated retrieval to extract alveolar space and vascular ROIs for targeted analysis. It then applied LMM analysis to evaluate extracellular matrix and immune population trajectories within each compartment.

The analysis revealed distinct dynamics within the two spatial compartments. Macrophage enrichment was observed as a pan-compartmental event occurring consistently across both alveolar and vascular spaces, while fibrotic scarring was strictly compartmentalized. Specifically, collagen deposition exhibited a highly significant, monotonic increase solely within the alveolar space (FDR *p* = 0.004). Conversely, its relative proportion remained flat in vessels (FDR *p* = 0.173). When evaluating alveolar neutrophils, *HypothesisValidator* detected high baseline variance across individuals (ICC = 60.8%). After adjusting for this intra-patient correlation, the LMM engine suppressed a region-level correlation (naive *p <* 0.001) and yielded a non-significant result (FDR *p* = 0.321). This demonstrates *OmicsNavigator*’s ability to mitigate the impact of pseudo-replication and isolate genuine, localized signals (e.g., alveolar fibrosis) while preventing the overestimation of statistical significance.

These results parallel and refine the findings of Rendeiro et al. [36]. Consistent with the original report of pan-tissue macrophage infiltration and progressive alveolar fibrosis in late COVID-19, *HypothesisValidator*’s compartment-decoupled analysis localized the collagen increase to the alveolar space—a finer-grained test not performed in the original study, where fibrosis was scored at the region level. Furthermore, *HypothesisValidator* used a patient-level mixed-effects framework to replace the original Mann–Whitney tests, which addressed the inter-individual variances better and avoided statistical overestimation due to pseudoreplications.

## 3 Discussion

The advent of highly multiplexed spatial omics has exposed an analytical bottleneck: the challenge of traversing large volumes of spatially resolved molecular data and extracting data-grounded biological insights. While the recent integration of LLMs has accelerated bioinformatics workflows by automating code execution and tool utilization, current approaches operate primarily as computational technicians that reproduce cell-centric analyses, and thus leave the interpretation gap unaddressed. Moreover, the unconstrained generative nature of LLMs risks the generation of false positive discoveries such as spurious correlations and statistically unsupported hypotheses. To address these limitations, we developed *OmicsNavigator*, an autonomous agentic system designed to act as an objective scientific partner for spatial biology.

*OmicsNavigator* utilizes a sequential framework composed of planning, interpretation, and analysis modules that mirrors the analytical workflow of domain experts. The system builds on the core ability to interpret micro-scale ROIs using autonomously curated biological prior knowledge and spatially aware, multi-modal observations. This is demonstrated and validated across diverse datasets spanning different kidney and lung diseases. Based on this fine-grained understanding of data, *OmicsNavigator* further employs a scalable semantic retrieval mechanism and an aggregation pipeline to discover, locate, and summarize disease-relevant structures from patient data, reconstructing patient-level disease profiles grounded in ROI-level observations. Together, these capabilities enable study-scale autonomous hypothesis validation, where *Omic-sNavigator* extracts and synthesizes visual and molecular evidence from micro-scale ROIs into concrete biological findings that are both data-grounded and statistically tested.

A defining design of *OmicsNavigator* is its structural commitment to objectivity. During development, we observed a concerning propensity for autonomous agents to engage in algorithmic *p*-hacking. When tasked with validating a user’s hypothesis without guardrails, agents sometimes exhibited suspicious behaviors such as shifting ROI inclusion boundaries or iteratively switching between parametric and non-parametric statistical tests to artificially achieve a significance threshold that satisfied the initial prompt. To prevent this, *OmicsNavigator* establishes mechanisms to enforce isolation of semantic reasoning/planning from statistical computations. By locking biological priors (i.e., Background Knowledge Checklists) and validation pipelines (i.e., Verification Blueprint) before any data evaluation, *OmicsNavigator* mitigates the operating agents’ ability to retrospectively alter targets or statistical testing choices. This pre-registration design helps to ensure an objective, reliable scientific partner for spatial biology research.

Beyond preventing algorithmic bias, *OmicsNavigator* adopts an auditable HITL framework to mitigate the “black-box” risks and to ensure alignment with domain expertise. Before large-scale computations begin, researchers are mandated to approve the generated Background Knowledge Checklists and Verification Blueprint, facilitated via a graphical interface (Supplementary Figure 3). This ensures a pre-execution consensus that mitigates both downstream hallucinations and user-induced confirmation bias. Complementing this, the DAG-based execution of hypothesis validation enables retrospective traceability for outcomes. *OmicsNavigator*’s underlying data logs maintain a mapping between any patient- or study-scale outcome (e.g., a reported *p*-value for a trend) and the exact ROI observations that contributed to it. Researchers can therefore unpack any high-level conclusion into a continuous, grounded chain of evidence down to the raw, micro-scale interpretations of individual ROIs. This enforced transparency of *OmicsNavigator* ensures that every autonomous insight remains fully verifiable.

Despite the demonstrated capabilities in automating spatially aware reasoning and hypothesis validation, *OmicsNavigator* operates within explicit technical boundaries. First, as a prior-constrained engine, its efficacy heavily depends on the quality of background knowledge. In “knowledge-sparse” scenarios—such as rare diseases or studies of highly heterogeneous tumor microenvironments—traditional API-based literature retrieval is inflexible and may miss critical mechanistic nuances. A potential improvement to overcome this bottleneck is to integrate “Deep Research” capabilities that execute multi-step navigation and comprehensive full-text parsing, synthesizing context-specific knowledge graphs. We envision that this addition will enhance the system’s adaptability to less-studied biological domains. Second, the reliance on proprietary, cloud-based LLMs introduces substantial operational bottlenecks. Profiling approximately 1,000 ROIs in our DKD cohort consumed roughly 3 million completion tokens (Supplementary Table B1), imposing financial barriers to large-scale studies. A practical optimization is a hybrid model routing architecture: retaining highly capable proprietary models for complex tasks (e.g., global planning and Deep Research), while migrating computationally intensive, high-throughput ROI interpretations to locally deployed, open-weight language models. This approach will substantially reduce operational costs while mitigating the reproducibility issues caused by silent backend updates of closed-source LLMs. Third, while the underlying agentic framework of *OmicsNavigator* can adapt to diverse data modalities, conducting joint reasoning across spatial multi-omics introduces more biological complexity and data alignment challenges. Future iterations may further enhance the Background Knowledge Checklists and interpretation modules to integrate information from multiple perspectives—gene expression, protein expression, metabolites—thereby cross-validating overlapping yet heterogeneous molecular layers. This will facilitate the construction of a more comprehensive and robust understanding of the tissue landscape.

The landscape of agentic scientific research is rapidly evolving with the emergence of highly capable, general-purpose systems (e.g., Claude Code). Leveraging these generic coding engines represents a promising trend in the further development of agentic scientific partners, as they demonstrate remarkable proficiency in planning and executing analysis pipelines when provided with structured data. In our experiments, directly deploying these generic agents in spatial omics studies still faces certain limitations, including hallucinations due to a lack of reliable prior biological knowledge and a pronounced bias towards using only the data tables while neglecting spatial contexts and visual morphology. From this perspective, the domain-specific designs in *OmicsNavigator* —reasoning guardrails via checklists and blueprint, spatially aware data interpretation and retrieval capabilities—provide complementary solutions. By packaging these capabilities into modular components—effectively serving as domain-specific “skills” or tool-calling functions for generic engines—developers can harness the best of both paradigms. This synergistic integration points toward the next generation of scientific tools: universally capable systems anchored and supported by domain-specific knowledge and skills.

As spatial technologies continue to map tissues at unprecedented scales, simply accumulating more data will not bridge the interpretation gap. Instead, we show in this study that agentic AI applications can enable spatially aware, objective data exploration and hypothesis validation at scale. This ongoing transition from using AI as computational technicians to AI as research partners will greatly expand the capabilities of human researchers and accelerate future discoveries in spatial biology.

## 4 Methods

### 4.1 Data Acquisition

#### 4.1.1 Diabetic kidney disease (DKD) dataset

We used a previously published CODEX dataset of kidney samples from diabetic patients [33], hereinafter referred to as the DKD dataset. A total of 17 regions comprising approximately 137,000 cells were used for *OmicsNavigator* experiments. These regions span multiple stages of diabetic kidney disease (early/DM to stage 3/DKD3) and exhibit a broad spectrum of disease-associated histological changes.

Each region in the DKD dataset was preprocessed using a standard pipeline involving cell segmentation, biomarker integration, and cell typing via Leiden clustering plus expert pathologist annotations, resulting in 11 identified cell types, including tubular, endothelial, immune, and other cells. Pathologist annotations of five major anatomical structures—proximal tubules, distal tubules, glomeruli, blood vessels, and interstitium—were used to evaluate *OmicsNavigator*’s zero-shot annotation accuracy in subsequent experiments. Clinical staging labels (DM, DKD2A, etc.) were used in the hypothesis validation case study to uncover stage-dependent trajectories of diabetic glomerulopathy.

#### 4.1.2 Transplant rejection (TR) dataset

We adapted a different kidney dataset [35] comprising samples profiled using CODEX from 5 normal and 7 transplant rejection kidney regions, hereinafter denoted as the TR dataset. In total, around 350,000 cells were identified and classified after the same preprocessing pipeline, yielding 20 different cell types including immune, tubular, fibroblast, and smooth muscle cells. The TR dataset was profiled using a different biomarker panel, with only 7 of the 52 markers (13%) overlapping with the DKD dataset.

#### 4.1.3 Kidney allograft rejection (KAR) dataset

Another dataset consisting of 8 T cell-mediated and 8 antibody-mediated rejection kidney cases [34] was used in parallel to test the generalizability of *OmicsNavigator*. This dataset similarly utilized a different biomarker panel and contained around 558,000 cells across 13 distinct cell types, with a particular focus on immune populations within kidney structures.

#### 4.1.4 COVID-19 and acute respiratory distress syndrome (ARDS) dataset

We utilized a previously published imaging mass cytometry (IMC) dataset of human pulmonary tissue from patients with different pathological conditions, including COVID-19 infection and ARDS [36]. The dataset targets the expression of 36 proteins to investigate the spatial architecture of acute lung injury, and to capture the temporal progression of the disease from early to late stages based on the time of symptom onset. The COVID-19 dataset contained a total of around 395,000 cells from 157 IMC images, collected from 14 patients. The analysis pipeline identified key alveolar epithelial cells, different immune infiltrates (including macrophages and neutrophils), and mesenchymal/fibroblast populations. Pathologist annotations of major anatomical structures—vessels, airway, alveolar space, etc.—were derived from a subsequent work [45] and used to evaluate *OmicsNavigator*’s zero-shot annotation accuracy in subsequent experiments. Clinical labels (healthy, early infection, late infection) were used in the hypothesis validation case study to examine pathogenesis of pulmonary SARS-CoV-2 infection.

Collectively, all four datasets utilized high-dimensional profiling techniques with customized biomarker panels using different technical platforms (immunofluorescence and mass spectrometry imaging). They encompassed different organ systems and disease phenotypes. Because cell segmentation and annotations were performed independently across the original studies, these pipelines yielded non-overlapping sets of cell types and features. *OmicsNavigator* operates on regions from each study in an independent, language-driven manner, thereby minimizing the influence of potential batch effects and panel discrepancies. Consequently, the interpretation and semantic search results generated by the system are directly comparable across studies.

### 4.2 System Architecture and Agentic Orchestration

*OmicsNavigator* operates as a multi-agent scientific partner. To mitigate the risk of circular reasoning and goal-drift typically observed in unconstrained, decentralized LLM networks, the cognitive workflow is orchestrated using a linear directed acyclic topology via the LangGraph framework. The system establishes a unidirectional chronological handoff across three isolated phases: Planning, Interpretation, and Analysis, **separated by a human-in-the-loop (HITL) audit checkpoint**. Inter-agent communication and memory management are standardized through a centralized, stateful dictionary (AgentState). Rather than employing dynamic message passing, this unified state acts as an immutable ledger that flows sequentially through the pipeline, continuously checkpointed in-memory.

*OmicsNavigator* is architected as a model-agnostic framework, decoupling the cognitive orchestration from any single proprietary LLM backend. To balance scientific creativity with analytical reproducibility across varied computational engines, *OmicsNavigator* implements a Stratified Cognitive Stochasticity Control strategy. For instance, when configured with the *gemini-2.5-pro* engine, exploratory agents (e.g., the *DataAnalyst* and *LiteratureReviewer*) operate under modulated thermodynamic parameters (temperature ∈ [0.2, 0.5]) to facilitate sufficiently broad and comprehensive contextual exploration. Conversely, downstream analytical agents tasked with structural interpretation and mathematical validation are clamped to determinism (temperature = 0.0, top p = 1.0), thereby reducing variance during quantitative data extraction and maintaining methodological reproducibility.

Long-term episodic memory for interpreted spatial structures is managed via a FAISS vector database. Natural language interpretations of Regions of Interest (ROIs) are vectorized utilizing the *gemini-embedding-001* model. The resulting high-dimensional semantic embeddings (*D* = 768) are evaluated using *L*_2_ distance metrics and immutably stored in FAISS. This architecture enables rapid, *O*(log *N*) semantic retrieval for subsequent population-level queries and spatial expansion algorithms, bypassing redundant LLM inference overhead during hypothesis validation.

### 4.3 Global Context Initialization

To ensure biological fidelity and restrict unconstrained LLM hallucinations during localized reasoning, *OmicsNavigator* initiates the analytical workflow by establishing a global cognitive context. This Plan Module operates prior to any microscopic region-of-interest (ROI) interpretation, decoupling biological prior formulation from downstream statistical execution.

The module deploys two specialized information-gathering agents. The *DataAnalyst* accesses raw spatial omics matrices via the FastMCP protocol to perform context-aware exploratory data analysis. Beyond computing foundational metrics (e.g., cellular composition, molecular dynamic ranges, and spatial entropy), the *Data-Analyst* operates within a **Closed-loop Statistical Reflection** mechanism. Upon generating an initial quantitative profile, the agent self-evaluates the mathematical validity and biological feasibility of the report. If critical gaps are identified, the agent autonomously formulates targeted analytical scripts, executes them within an isolated Python sandbox, and integrates the findings. This iterative self-correction loop ensures the baseline metrics are dynamically tailored to the specific spatial heterogeneity of the target sample.

Concurrently, the *LiteratureReviewer* retrieves verified patho-physiological priors. Rather than executing unconstrained programmatic searches, this agent parses the data profile to extract core metadata (e.g., highly expressed biomarker lists, unique cell types) and populates a highly structured semantic query template encompassing critical dimensions such as target microenvironments and expected mechanistic events. To transition from cognitive formulation to knowledge extraction, the *LiteratureReviewer* leverages native agentic web-browsing capabilities to execute targeted natural language queries derived from this template. By autonomously navigating and parsing primary evidence from authoritative biological repositories (e.g., NCBI, PubMed, and QuickGO), the agent synthesizes the extracted patho-physiological mechanisms and biomarker metadata into a comprehensive report. This execution logic ensures that the subsequently generated biological constraints are grounded in context-aware, peer-reviewed scientific literature rather than the unverified parametric memory of the LLM.

The gathered global statistics and literature priors are synthesized by the central *Planner* agent to orchestrate all downstream interactions. A critical output of this synthesis is the generation of a comprehensive domain-specific heuristic manual, instantiated as four distinct constraint checklists (VisualProfiler checklist.md, OmicsProfiler checklist.md, Disease Relevance Score Rubrics Checklist, and OmicsInterpreter checklist.md). These documents inject biological priors into the downstream Interpretation Module—for instance, explicitly defining channel-specific visual cues or establishing conflict resolution typologies (e.g., prioritizing morphological degradation over static molecular alterations in early-stage lesions). This ensures cross-modal discrepancies are resolved using grounded patho-physiological logic.

Furthermore, to enforce statistical rigor during hypothesis validation, the *Planner* generates a Verification Blueprint.json. At this critical juncture, *OmicsNavigator* suspends autonomous execution to enforce a HITL pre-analytical audit. Researchers are presented with the generated heuristic checklists and the blueprint to systematically review the intended analytical workflow. This “Plan Mode” architecture allows domain experts to verify the extracted biological priors, inject supplementary constraints, and correct potential misalignments before any computationally expensive procedures occur. Upon receiving explicit human confirmation, the blueprint transitions into a pre-registration document. Before any quantitative ROI evaluation commences, it locks the target statistical variables, cohort inclusion/exclusion criteria, and the declarative topology of the statistical testing DAG.

### 4.4 Multi-modal Interpretation Engine

At the core of *OmicsNavigator* is the Multi-modal Interpretation Engine, a sequential pipeline designed to process selected ROIs by synergizing morphological features with high-dimensional molecular signatures.

#### 4.4.1 VisualProfiler

The *VisualProfiler* is tasked with extracting morphological features and structural inferences from imaging data (e.g., multiplexed immunofluorescence). To provide the vision-language model with both precise cellular details and broader spatial context, we employ a *Dual-scale Visual Prompting* strategy. For each Anchor ROI, the agent receives two distinct image inputs:

1. A macroscopic contextual field-of-view (FOV), generated by downsampling the raw multi-channel image to a 512 × 512 pixel resolution, with the target ROI delineated by a bounding box to reveal its immediate tissue neighborhood.
2. A high-resolution, 512 × 512 pixel zoomed-in crop of the exact ROI boundaries.

Guided by the explicitly injected VisualProfiler checklist.md (which dictates channel-specific visual cues), this dual-input strategy ensures the model accurately infers complex anatomical structures while contextualizing local morphology within the global tissue architecture.

#### 4.4.2 OmicsProfiler

The *OmicsProfiler* translates quantitative, high-dimensional molecular matrices into discrete natural language representations via an Omics-to-Text Encoding algorithm. To establish a spatially-weighted baseline expression profile resilient to variations in cellular density, *OmicsNavigator* constructs a reference distribution through a three-step stochastic spatial sampling algorithm:

1. **Global Sliding Window Partitioning:** The entire spatial region is exhaustively partitioned using a sliding grid approach (patch size = 128 × 128 pixels, stride ratio = 0.5).
2. **Density Filtering and Hard Capping:** Sparse patches containing fewer than *n*_min_ = 5 cells are discarded. For dense patches exceeding *n*_max_ = 50 cells, the system enforces a hard cap by uniformly sampling exactly 50 cells. This step guarantees that the contribution of any microenvironment to the global baseline is proportional to its physical spatial area rather than its absolute cellular abundance.
3. **Stochastic Reference Assembly:** From the filtered candidate pool, *N* = 1000 patches are stochastically sampled without replacement. The aggregated samples constitute the robust Reference Subset.

For a target ROI *i* and biomarker *g*, let *x_i,g_*denote the initial normalized expression. The spatially-weighted background context is established by calculating the mean *µ_g_* and standard deviation *σ_g_* across the Reference Subset. To handle the extreme sparsity and zero-variance artifacts inherent to spatial omics data (where a rare biomarker might exhibit *σ_g_* = 0 within the reference pool), we implement a regularized generalized Z-score defined as:

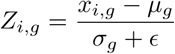

where *ɛ* is a positive regularization constant (set to *ɛ* = 10^−8^).

To categorize biomarker expression, *OmicsNavigator* employs a composite thresholding strategy. Guided by the OmicsProfiler checklist.md, a biomarker *g* is assigned semantic descriptors according to the following logic:

- **Highly Expressed:** The biomarker satisfies at least one of the following: local *Z_i,g_ >* 1.5, expression above the 90th percentile of the Reference Subset distribution, or it ranks among the top 2 highest expressed markers in the target cell.
- **Lowly Expressed:** The biomarker is not categorized as highly expressed, and satisfies either *Z_i,g_ <* −1.5 or falls below the 10th percentile of the Reference Subset distribution.
- **Moderately Expressed:** Biomarkers not meeting the aforementioned thresholds.

#### 4.4.3 OmicsInterpreter

The *OmicsInterpreter* synthesizes the visual inferences from the *VisualProfiler* and the textual omics matrices from the *OmicsProfiler* using a context-aware Late Fusion paradigm. Rather than executing unconstrained logical deductions, the interpreter’s cognitive space is governed by the OmicsInterpreter checklist.md generated during the initial Plan Phase. Guided by these grounded biological priors, the agent resolves cross-modal conflicts for the Anchor ROIs, generating a structured JSON report containing the inferred identity and a cognitive confidence score.

Concurrently, to facilitate downstream patient-level disease trajectory reconstruction, the *OmicsInterpreter* quantifies a localized Disease Relevance Score (ranging from 0 to 100). This continuous severity metric is dynamically governed by a Disease Relevance Score Rubrics Checklist. Ultimately, the inferred identity, confidence score, and disease relevance metric are vectorized and stored within the FAISS database, completing the episodic memory integration.

### 4.5 Scalable Retrieval via Cluster-Anchor-Expand

To overcome the computational bottleneck of querying the LLM for every individual ROI across gigapixel tissue landscapes, *OmicsNavigator* employs a highly scalable “Cluster-Anchor-Expand” algorithmic framework. This approach distills cohort-scale landscapes into a minimal set of representative Anchor ROIs for deep semantic interpretation, followed by dynamic spatial expansion during the retrieval phase.

#### 4.5.1 Multi-modal Featurization and Robust Scaling

To manage the memory constraints associated with Whole Slide Images (WSIs) and mitigate localized batch effects, the tissue landscape is initially partitioned into distinct geographic shards (instantiated as RegionShardData). Within each shard, for every ROI, the system constructs a concatenated multi-modal feature vector *v* ∈ R*^D^*that encapsulates both morphological and molecular states. The vector is derived from four independent feature matrices: image-level intensity histograms (*F*_img_), mean cellular biomarker expression (*F*_bm_), normalized cell-type frequency distributions (*F*_type_), and mean cellular morphological metrics (*F*_morph_).

To address dimensional and scaling discrepancies across these heterogeneous modalities, we apply a normalization protocol prior to concatenation. For each modality *m* ∈ {img, bm, type, morph}, let *D_m_* denote its pairwise Euclidean distance matrix across all ROIs within the shard. The normalized feature matrix *F’_m_* is computed as:

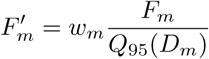

where *w_m_* serves as a predefined modality weight, and *Q*_95_(*D_m_*) represents the 95th percentile of the upper triangular elements of *D_m_*. This quantile-based scaling factor harmonizes global variance across modalities while preserving robustness against extreme biological outliers. The final representation for each ROI is the concatenated vector 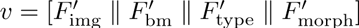.

##### Spatial Geometric Stratification

To ensure that the subsequently sampled Anchor ROIs are geographically dispersed across the tissue landscape rather than heavily localized within a single dense region, a secondary spatial partitioning step is introduced. Specifically, a *K*-Means algorithm is applied explicitly to the *xy*-spatial coordinates of the ROIs within each primary phenotypic cluster. The number of geometric partitions (*K*_spatial_) is adaptively parameterized based on the physical dimensions of the target tissue shard (e.g., *K*_spatial_ = 3 for standard regions). While this centroid-based approach does not delineate exact, density-based anatomical boundaries, it acts as a geographic stratifier—segmenting the widely distributed phenotypic cluster into distinct geometric sectors (i.e., spatial quadrants). This spatial stratification forces the downstream stochastic sampling mechanism to draw representative anchors from diverse physical territories of the tissue shard, thereby mitigating regional sampling bias.

#### 4.5.2 Distance-Weighted Stochastic Anchor Sampling

To select representative Anchor ROIs for LLM interpretation, we eschew nearest-centroid selection to avoid sampling unrepresentative edge cases. Instead, we implement a distance-weighted stochastic sampling mechanism coupled with dense-core truncation.

For a given sub-cluster containing *N* ROIs, let *ṽ* denote the component-wise median feature vector. The phenotypic Euclidean distance for the *i*-th ROI is calculated as *d_i_* = ‖*v_i_* − *ṽ*‖_2_. An initial inverse-distance probability vector *p* is formulated as 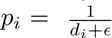.

To restrict sampling to the core phenotypic identity of the cluster, we apply a hard median truncation, zeroing out the selection probability of any ROI falling outside the top 50% of phenotypic proximity to the median. The truncated probability 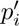 is defined as:

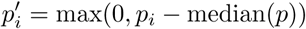

Finally, a non-linear probability sharpening step is applied via squaring 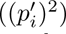 to disproportionately amplify the sampling likelihood of ROIs situated closest to the phenotypic core. The ultimate sampling probability *P* (*i*) for the *i*-th ROI is defined by the following normalization:

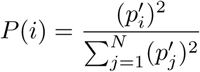

Using this distribution, a target number of Anchor ROIs (*k*) is sampled without replacement from each sub-cluster. This sampling depth is parameterized as an adaptable hyperparameter (e.g., *k* = 2 is utilized for the standard tissue cohorts evaluated in this study).

#### 4.5.3 Dynamic Expand-on-Hit Retrieval

These sampled Anchor ROIs are the targets dispatched to the Multi-modal Interpretation Engine, and only their resultant semantic interpretations are vectorized into the FAISS database. *OmicsNavigator* bypasses premature, global label propagation. Instead, the “Expand” phase operates dynamically within the *SemanticRetriever* agent during the querying phase.

When a user’s natural language query semantically matches a specific Anchor ROI within the FAISS index, the system executes an “Expand-on-hit” operation. It utilizes the SpatialPivotRegistry to trace the hit back to its original sub-cluster and conditionally propagates the relevance score to its geographic members. This lazy-evaluation architecture enables robust patient-level statistical aggregations without incurring the prohibitive token costs or vector database bloat associated with exhaustively computing LLM interpretations for hundreds of thousands of individual cells.

### 4.6 Statistical Hypothesis Validation

To balance the necessity of dynamically adapting to diverse spatial omics experimental designs with strict objectivity, *OmicsNavigator* employs a decoupled hypothesis validation engine. The architecture separates analytical strategy formulation from mathematical execution and final semantic synthesis, thereby harnessing the reasoning capabilities of LLMs while mitigating algorithmic *p*-hacking.

#### Autonomous Analytical Planning and Pre-registration

During the initial Plan Phase, the *Planner* agent autonomously architects a bespoke statistical strategy tailored to the specific study. To prevent algorithmic data-snooping, the *Planner* is isolated from the raw expression matrices and relies on the quantitative profile and baseline metrics generated by the upstream *DataAnalyst*. By parsing this context-aware exploratory report, the *Planner* evaluates the global structural topology and clinical metadata—such as cohort scale, hierarchical nesting (e.g., multiple regions sampled per patient), and the nature of clinical variables. Based on these architectural properties, the *Planner* synthesizes its comprehensive knowledge of scientific computing libraries (e.g., scipy.stats, statsmodels) to autonomously define the optimal mathematical modeling approach. For instance, upon detecting high technical-to-biological replicate ratios, it formulates Linear Mixed Models (LMM, e.g., *Feature* ∼ *Stage*+(1|*Patient ID*)) to account for intra-patient variance and quantify Intraclass Correlation (ICC). This logic, alongside explicitly logged fallback edges and False Discovery Rate (FDR) parameters, is structured into a formal Directed Acyclic Graph (DAG) and serialized into the Verification Blueprint.json. Following an HITL audit, this blueprint transitions into a pre-registration document, locking the statistical methodology before any quantitative hypothesis evaluation commences.

#### Execution and Context-Aware Synthesis

In the validation phase, a computational node traverses the pre-registered DAG, executing the prescribed formulas and applying global FDR control (via the Benjamini-Hochberg procedure) to all terminal *p*-values. Upon completion, the system re-engages the reasoning capabilities of the *HypothesisValidator* agent. The statistical facts—comprising FDR-corrected *p*-values, ICCs, and trajectory coefficients—are passed to the agent as a factual matrix. Operating as an intelligent evaluator, the *HypothesisValidator* analyzes these mathematical results within the broader spatial and biological context. It synthesizes these facts to formulate the final scientific evaluation (e.g., VERIFIED or REFUTED), extracting nuanced insights such as contextualizing high spatial variance (high ICC) as patient-specific resilience, or interpreting misaligned coefficient peaks as evidence of an asynchronous biological cascade. Because the agent cannot alter the upstream execution DAG or the resulting metrics, its autonomy is confined to contextual interpretation grounded strictly in verified data. The final output is a comprehensive natural language summary (validation report.md) that explicitly couples the agent’s contextual reasoning side-by-side with the raw statistical evidence, ensuring complete transparency for human expert review.

### 4.7 Evaluation Metrics

To assess the performance of OmicsNavigator across both zero-shot interpretation and semantic retrieval tasks, we established ground truth human annotations aggregated from the original cell-level metadata of the respective datasets.

#### 4.7.1 Zero-Shot Annotation Accuracy

For macroscopic structural interpretation, performance was evaluated utilizing the Macro *F*_1_-score to ensure minority phenotypic classes were equally represented. Given that the underlying datasets inherently provide annotations at the single-cell level, the structural ground truth for any evaluated Region of Interest (ROI) was established via a plurality vote. Consequently, the ROI-level semantic prediction generated by *Omic-sNavigator* was recorded as a True Positive when it perfectly matched this aggregated ROI ground truth.

The Macro *F*_1_-score is defined as the unweighted arithmetic mean of the class-specific *F*_1_-scores across all valid semantic classes *C*:

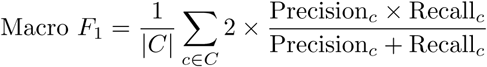

where Precision*_c_*and Recall*_c_*correspond to the precision and recall computed independently for each tissue architecture class *c*.

#### 4.7.2 Semantic Retrieval Performance

To evaluate the system’s capacity to accurately rank and retrieve phenotypically relevant microenvironments based on natural language queries, we utilized the Mean Average Precision at *K* (*mAP* @*K*). A retrieved ROI was classified as a relevant True Positive hit if its aggregated ground-truth label aligned with the target semantic intent of the query.

The metric is defined as the mean of the Average Precision scores across all queries:

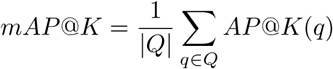

where *Q* represents the exhaustive set of standardized semantic queries, and *K* denotes the predefined rank cutoff for the retrieval list. The Average Precision at *K* for a specific query *q*, denoted as *AP* @*K*(*q*), is explicitly formulated as:

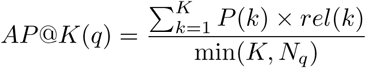

In this formulation, *P* (*k*) is the precision computed at cutoff rank *k*, *rel*(*k*) is an indicator function equating to 1 if the item at rank *k* is a relevant True Positive and 0 otherwise, and *N_q_* denotes the total number of actual relevant ROIs for query *q* existing within the entire database.

### 4.8 Code and Data Availability

The code and data of OmicsNavigator are available at https://github.com/yyli-leo/ OmicsNavigator.

Public datasets used in this study can be accessed at:

- Kidney studies: https://zenodo.org/records/12682727
- Healthy lung tissues: https://zenodo.org/records/6376767
- SARS-CoV-2 infected lung tissues: https://zenodo.org/records/4110560

## Appendix A Supplementary Figures

**Supplementary Fig. 1.**
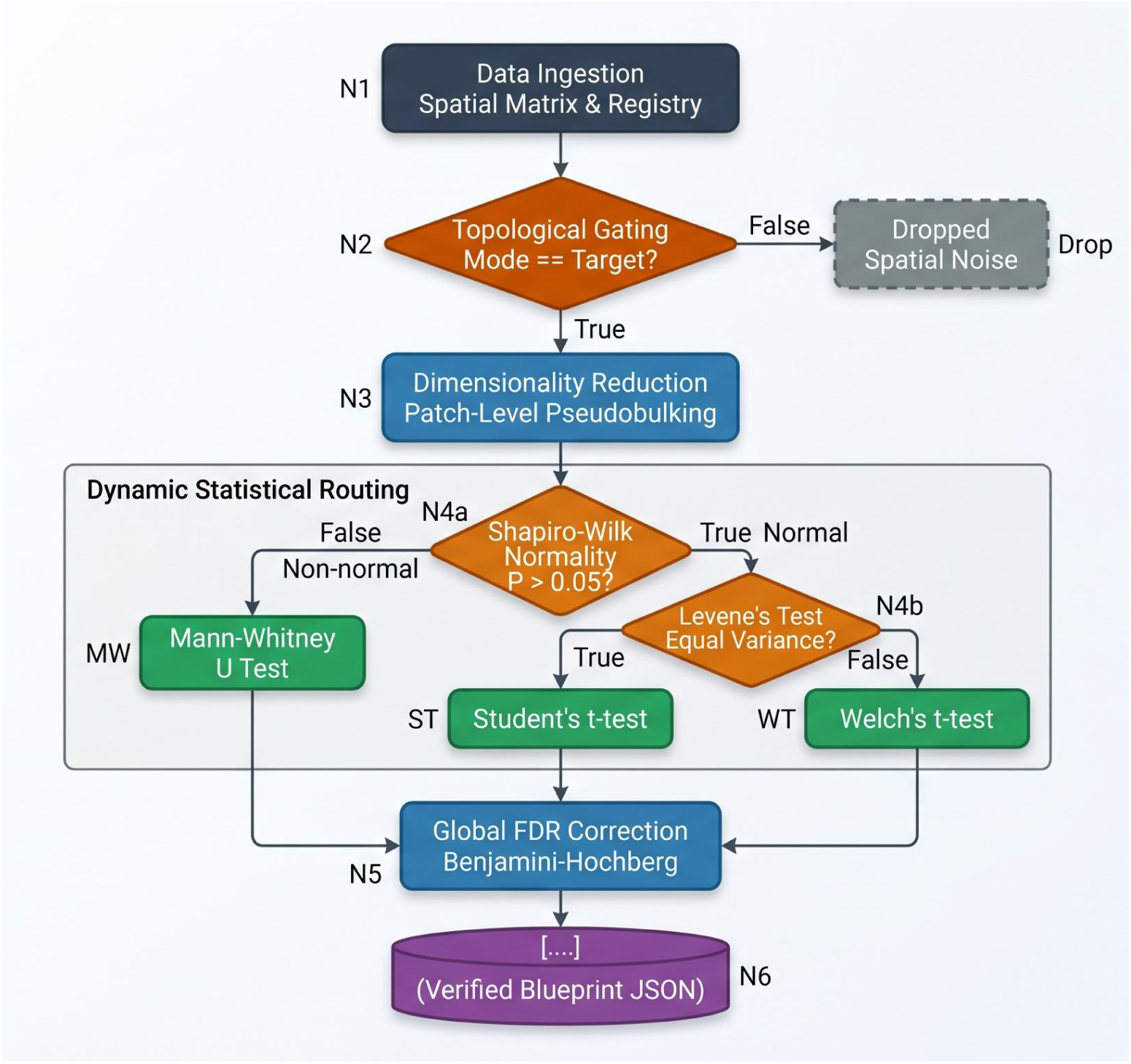
Deterministic execution pipeline via Directed Acyclic Graph (DAG) for autonomous hypothesis validation. This schematic visualizes a representative hard-coded mathematical topology governing the *HypothesisValidator* agent. The execution logic is predefined and constrained by the pre-registered Verification Blueprint (Supplementary Note B.3). Upon data ingestion (N1), the system filters Regions of Interest based on pre-registered topological criteria (N2) and performs pseudobulking (N3). The statistical routing (N4) dynamically routes comparative data through a decision tree guided by objective assumption checks: Shapiro-Wilk test for normality (N4a) and Levene’s test for homoscedasticity (N4b). Based on these Boolean outcomes, data are evaluated via Mann-Whitney U test, Student’s t-test, or Welch’s t-test, followed by a Benjamini-Hochberg False Discovery Rate (FDR) correction (N5). The final empirical metrics are structured into a read-only verified JSON output (N6).

**Supplementary Fig. 2.**
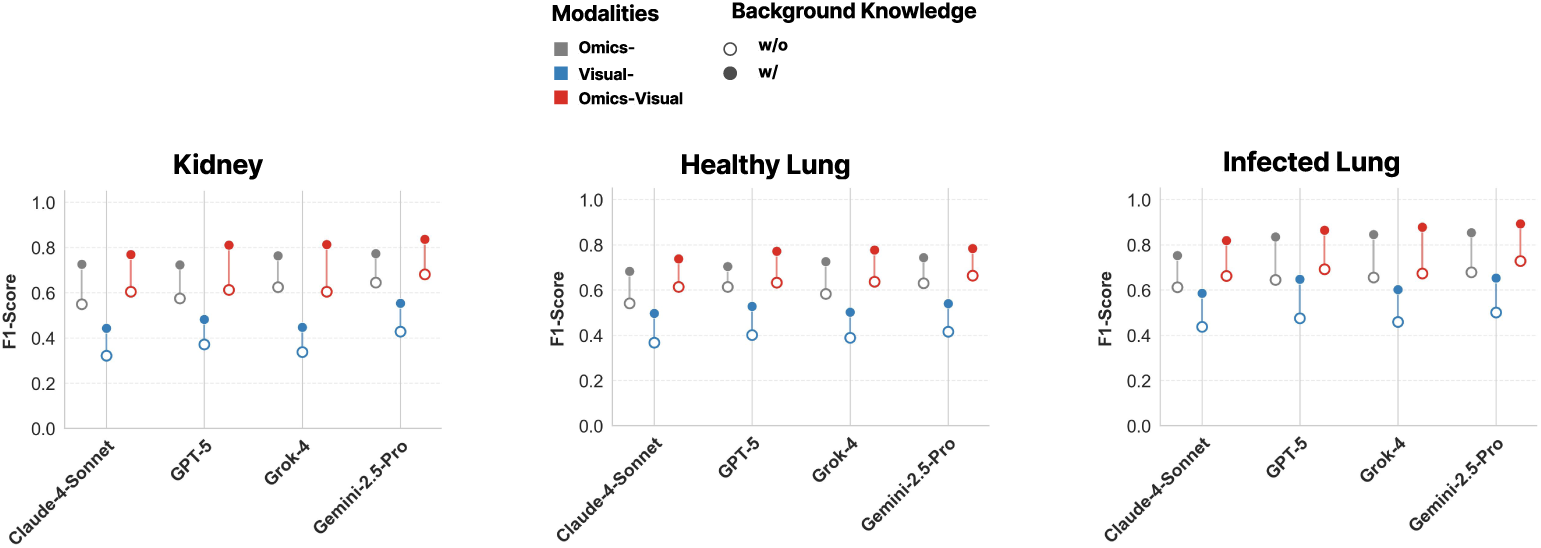
Ablation analysis demonstrating the critical impact of autonomously generated biological priors on interpretation accuracy. Dumbbell plots illustrating the zero-shot structural annotation performance (Macro F1-score) across three distinct tissue datasets: Kidney (left), Healthy Lung (center), and COVID-19 Infected Lung (right). Performance is systematically benchmarked across four state-of-the-art foundational models (Claude-4-Sonnet, GPT-5, Grok-4, and Gemini-2.5-Pro). Colors denote the input modality: Omics-only (grey), Visual-only (blue), and Omics-Visual fusion (red). Open circles (*◦*) indicate the baseline baseline performance of unconstrained models without background knowledge. Solid circles (*•*) represent the performance of models guided by the autonomously generated Background Knowledge Checklists. The vertical connecting lines visualize the performance gain achieved through knowledge injection. The results reveal a universal and substantial improvement in annotation accuracy across all data modalities, model architectures, and biological contexts when domain-specific priors are introduced. Notably, the complete *OmicsNavigator* configuration (multi-modal fusion with background knowledge; solid red circles) consistently yields the highest F1 scores, underscoring the necessity of the Plan Module in anchoring agentic reasoning and mitigating LLM hallucinations.

**Supplementary Fig. 3.**
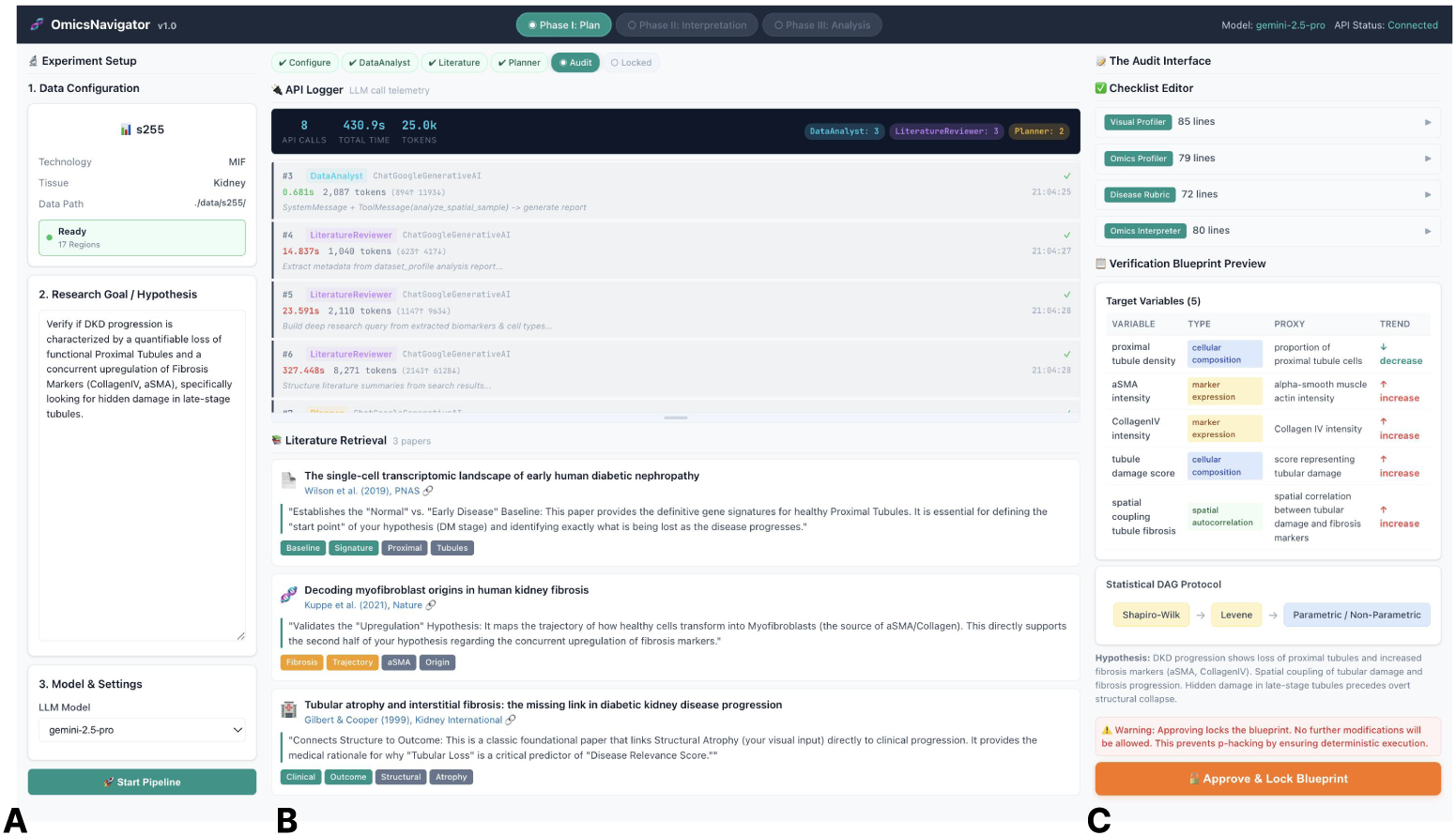
The Human-in-the-Loop (HITL) graphical interface for pre-execution audit and hypothesis pre-registration. The *OmicsNavigator* user interface (UI) is designed to enforce transparency, mitigate black-box risks, and establish analytical guardrails prior to large-scale computation. The interface consists of three functional panels during Phase I (Plan Module). (a) Configuration and State Monitoring (Left Panel). Researchers define the spatial data paths, select the underlying LLM architecture, and input the natural language hypothesis. (b) Agentic Reasoning and Literature Retrieval (Center Panel). A real-time API Logger tracks the token consumption, latency, and tool-use behaviors of the *DataAnalyst* and *LiteratureReviewer* agents. The retrieved, context-aware biological literature backing the derived heuristics is displayed with direct links. (c) The Audit Interface and Pre-registration (Right Panel). Before any micro-scale ROI interpretation or statistical analysis commences, the human researcher is mandated to review the autonomously generated outputs. This includes the four editable Background Knowledge Checklists (top right) and the Verification Blueprint (bottom right). The blueprint transparently displays the locked target variables, expected biological trends, and the deterministic Directed Acyclic Graph (DAG) statistical routing. By explicitly clicking the “Approve & Lock Blueprint” button, the researcher locks the configurations, transitioning them into read-only, pre-registered documents.

## Appendix B Supplementary Notes

### B.1 Computational efficiency and resource consumption

**Supplementary Table B1.** Resource Consumption and Cost Estimation of OmicsNavigator. Performance metrics are benchmarks conducted on the Diabetic Kidney Disease (DKD) dataset using the Gemini-2.5-Pro engine. Cost estimations are calculated based on the specific context window and caching pricing models at the time of execution. Note that metrics for the Interpretation Module represent the cost associated with a single Region of Interest (ROI).

| Phase | Token Usage |  |  | API Latency (s) | Estimated Cost (USD) |
| --- | --- | --- | --- | --- | --- |
|  | Uncached Input | Cached Input | Completion |  |  |
| Plan Module | 9000 | 0 | 28000 | 800 | \$0.29 |
| Interpretation Module<sup>a</sup> | 2000 | 4500 | 3000 | 46 | \$0.05 |
| Analysis Module | 2500 | 0 | 6000 | 60 | \$0.06 |

<sup>a</sup> Metrics for the Interpretation Module are reported **per ROI**. Total session cost scales linearly with the number of queried ROIs.
*Note:* Cost estimations are based on Gemini-2.5-Pro pricing: Input \$1.25–2.50/1M tokens, Output \$10.00–15.00/1M tokens, and Context Caching \$0.125–0.25/1M tokens, depending on context length (< or > 200k tokens).

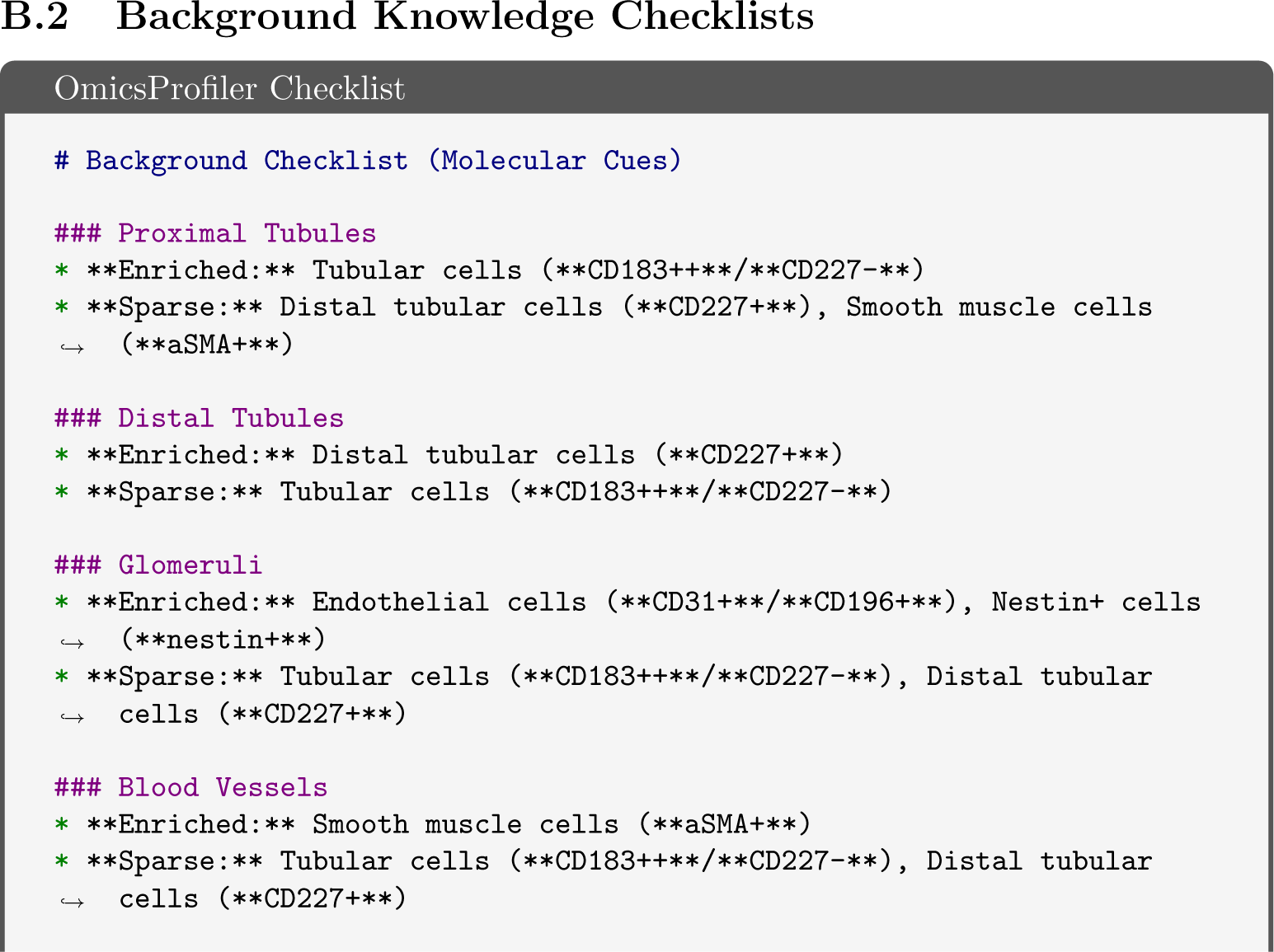
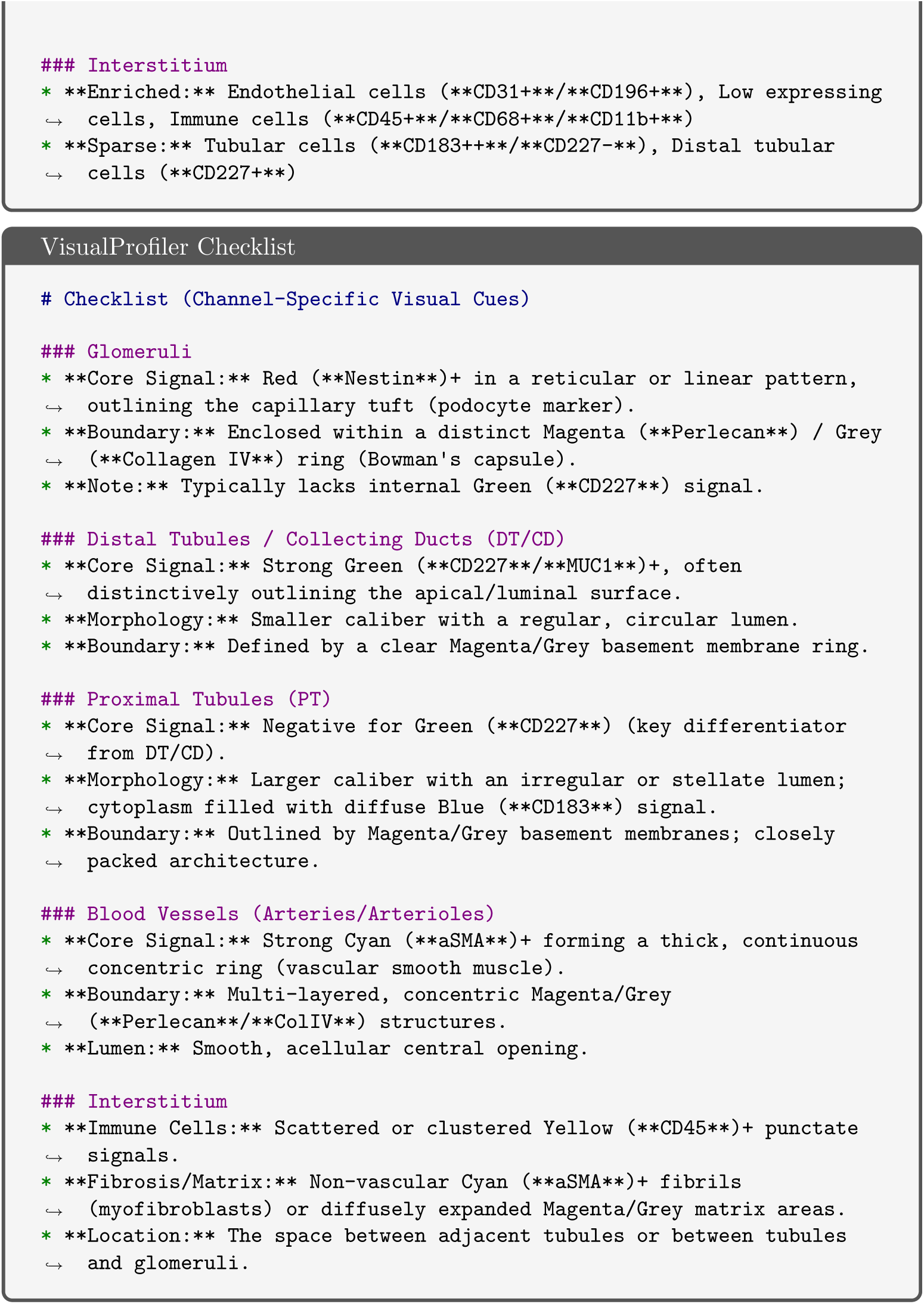
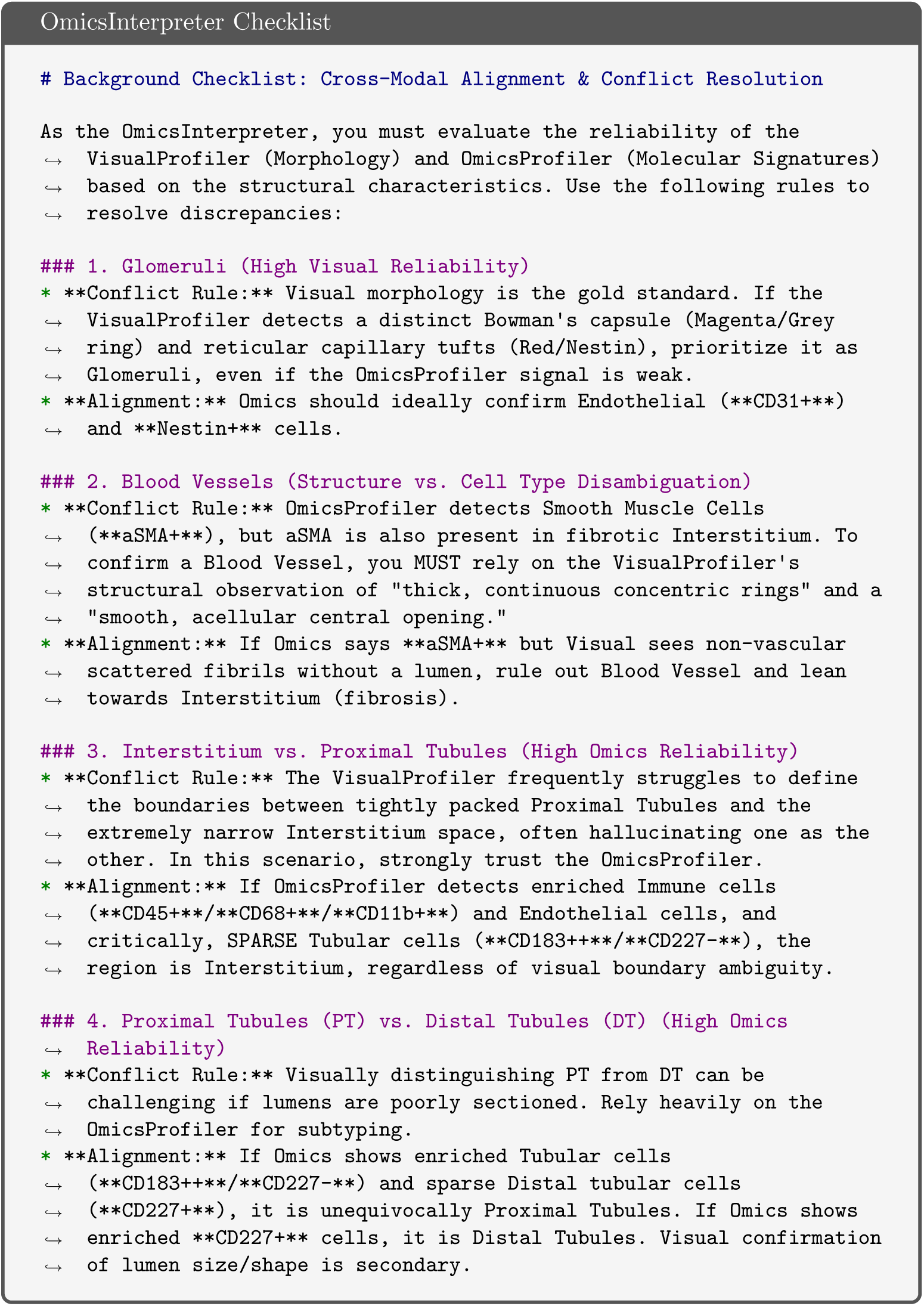
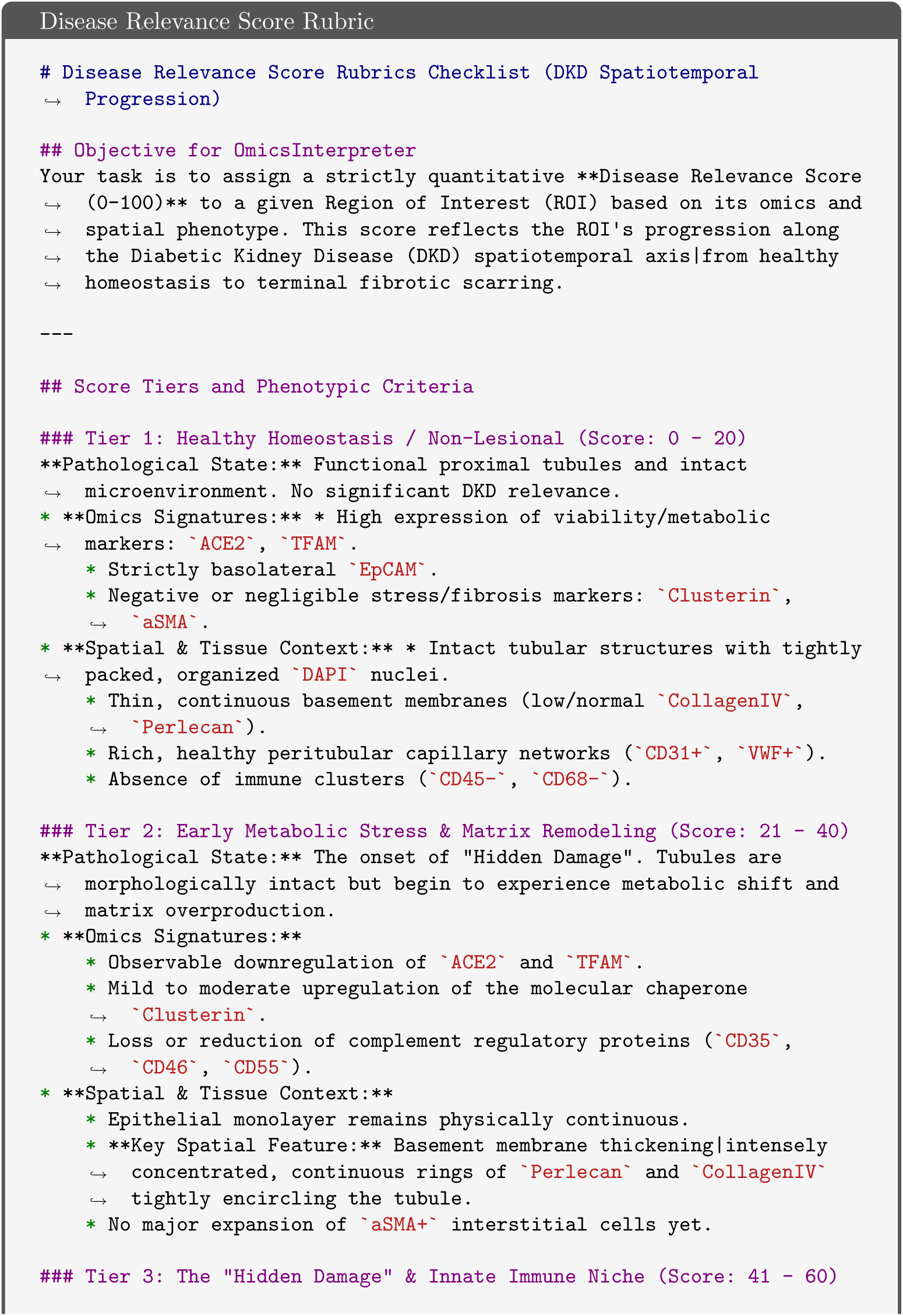
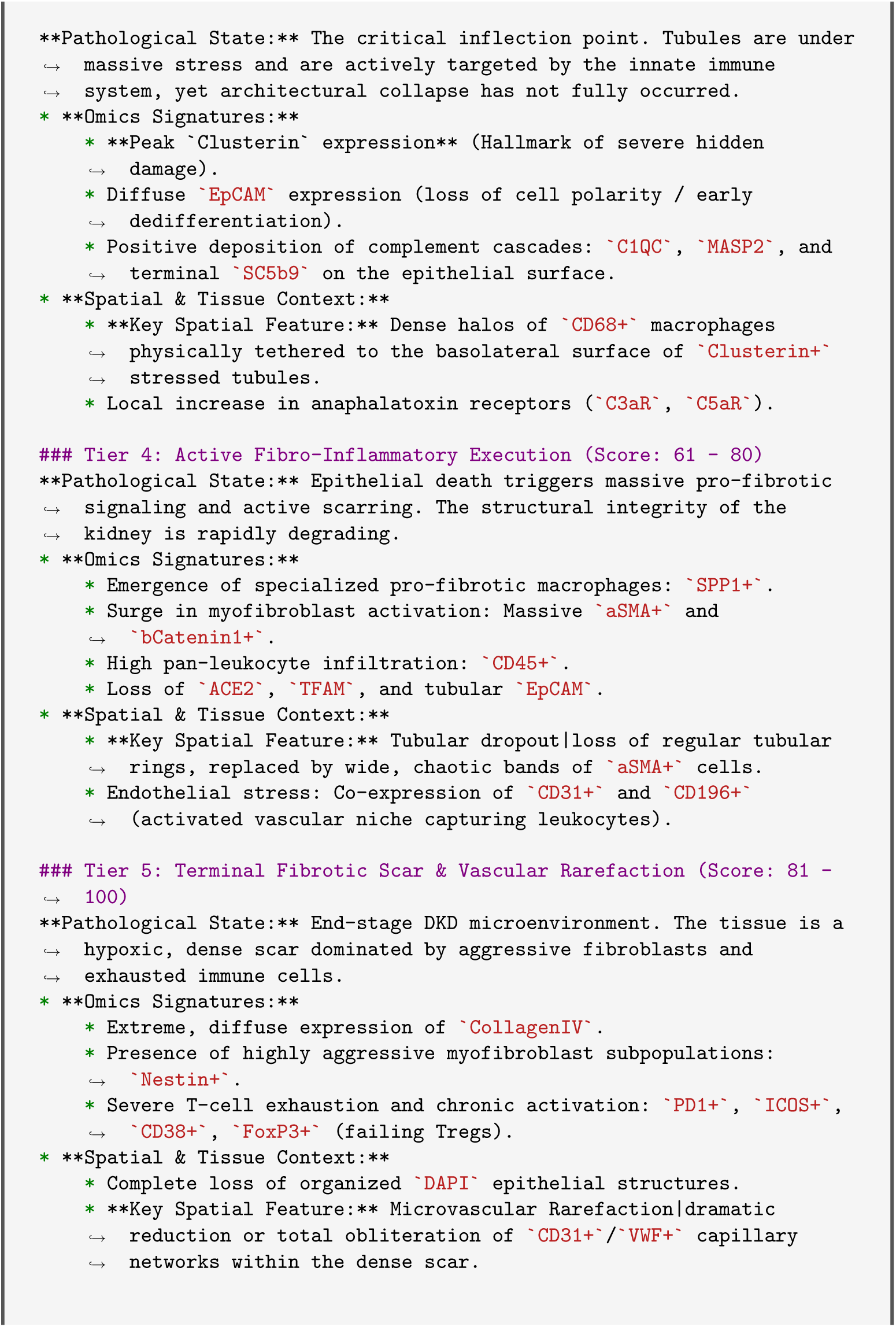
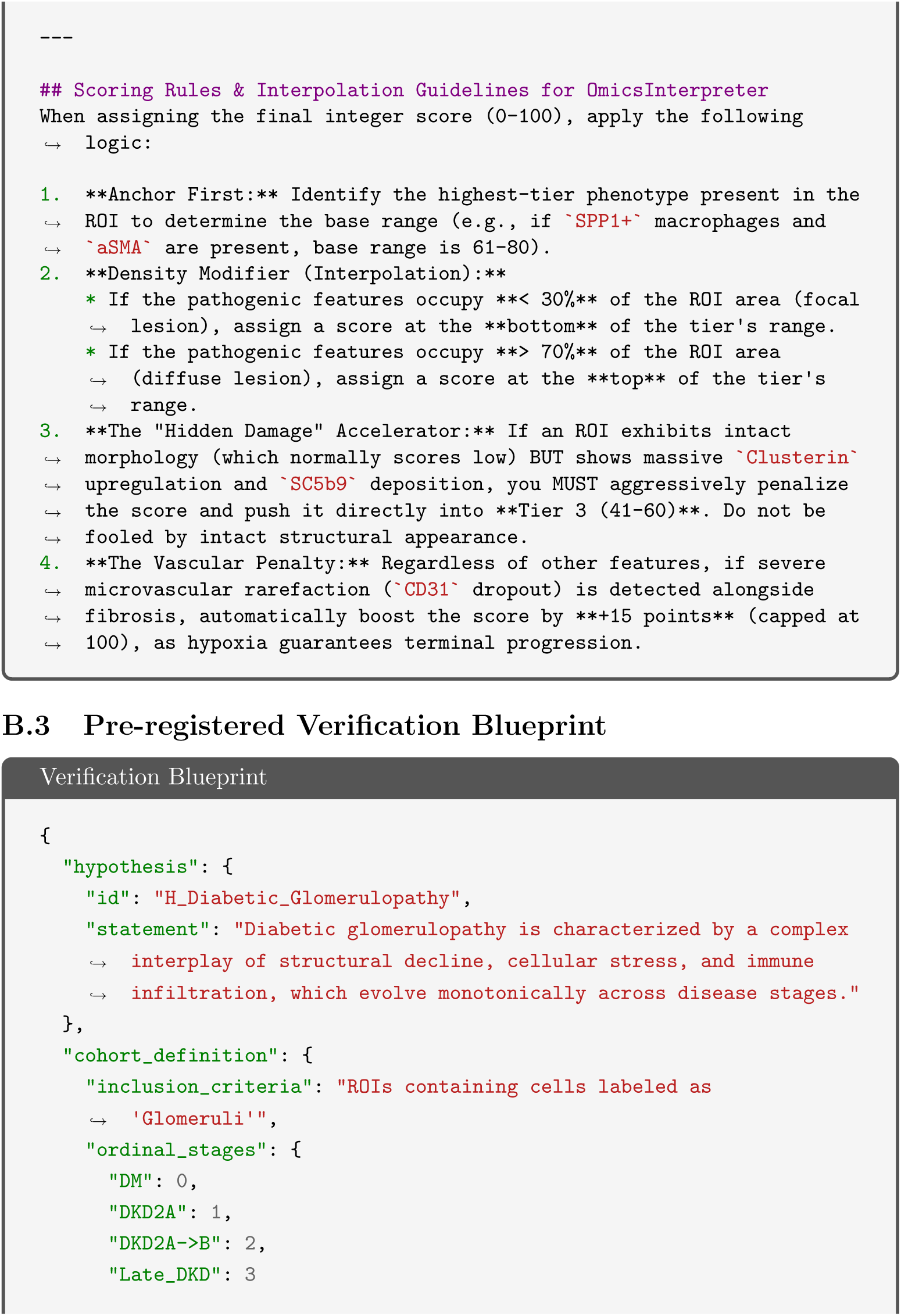
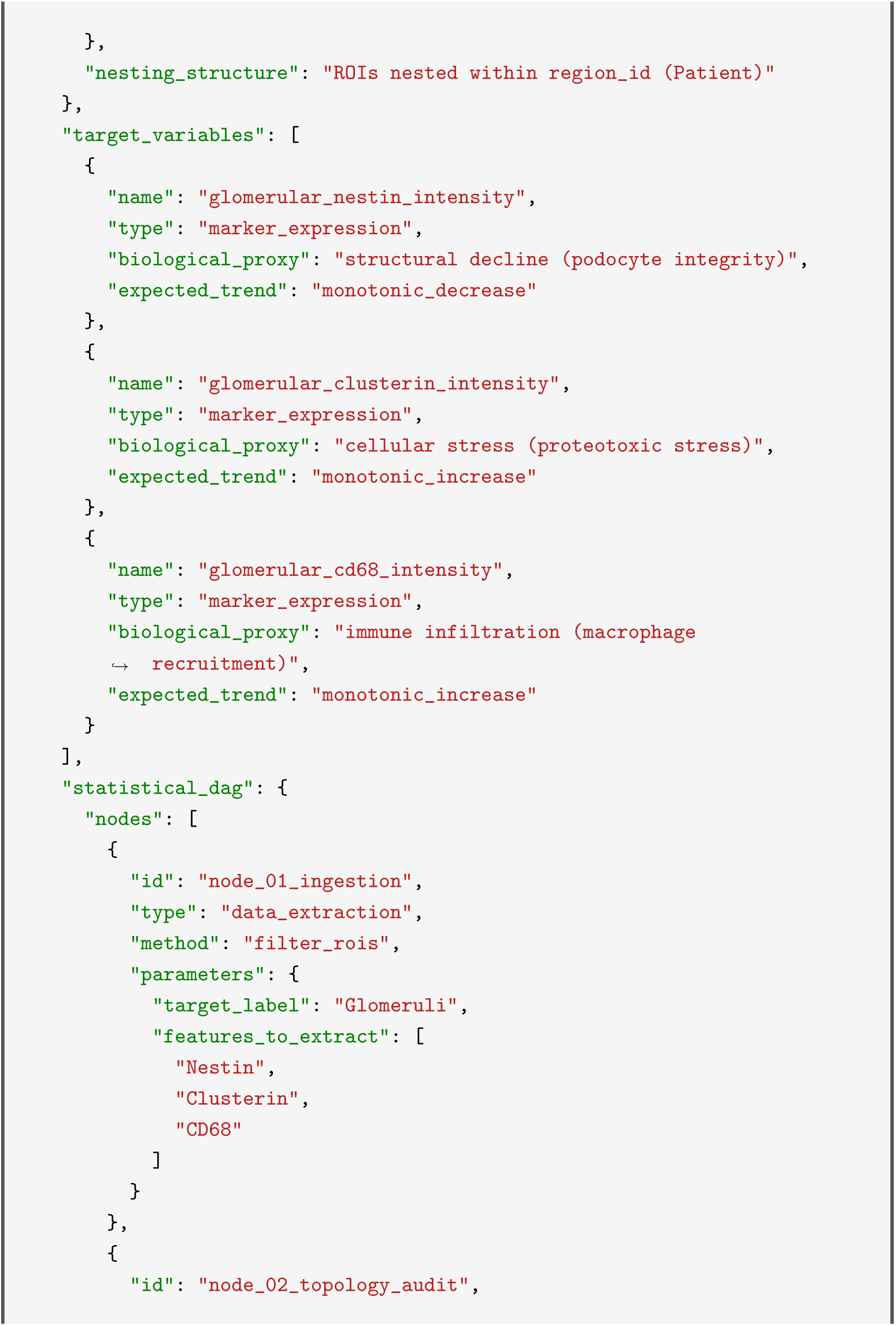
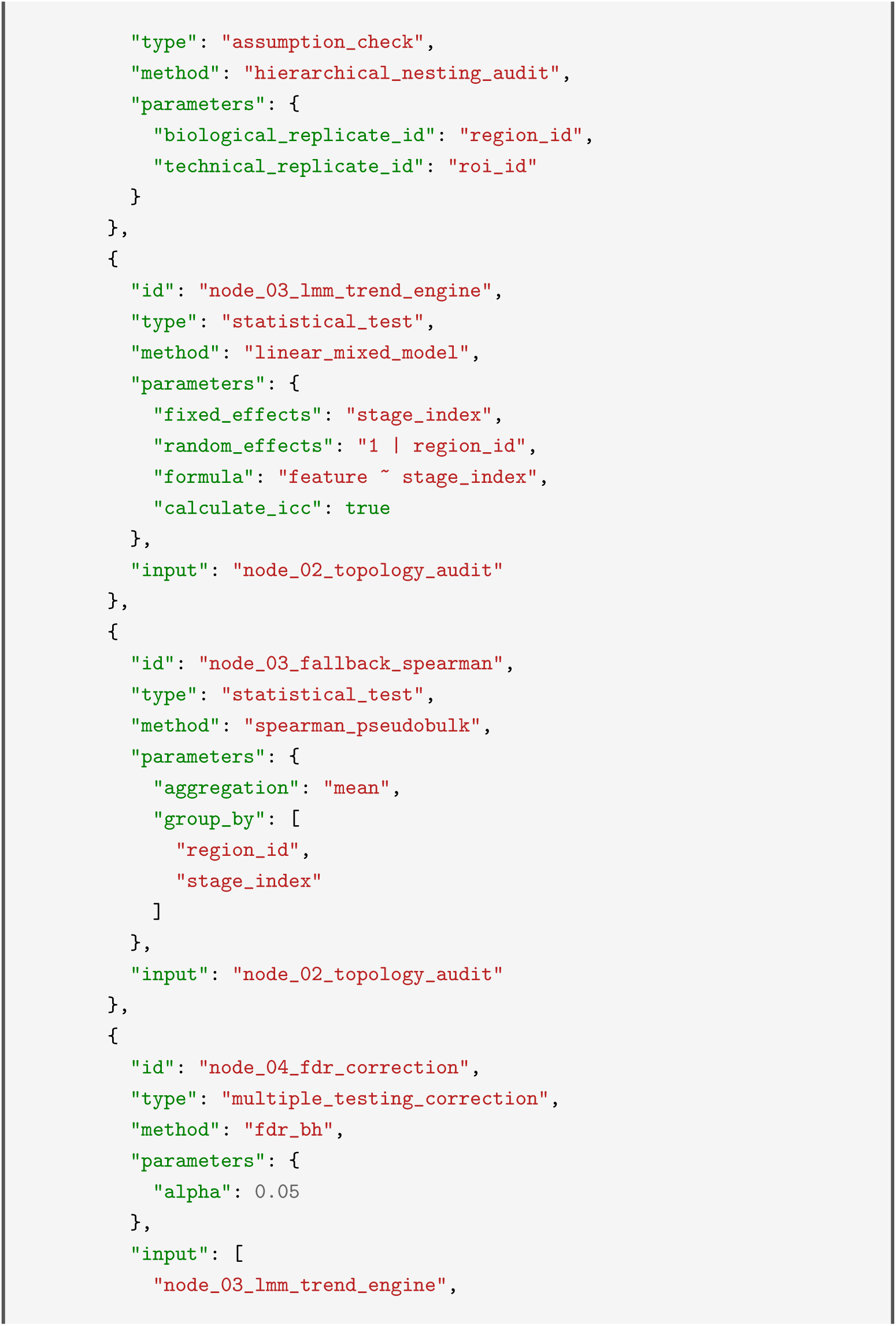
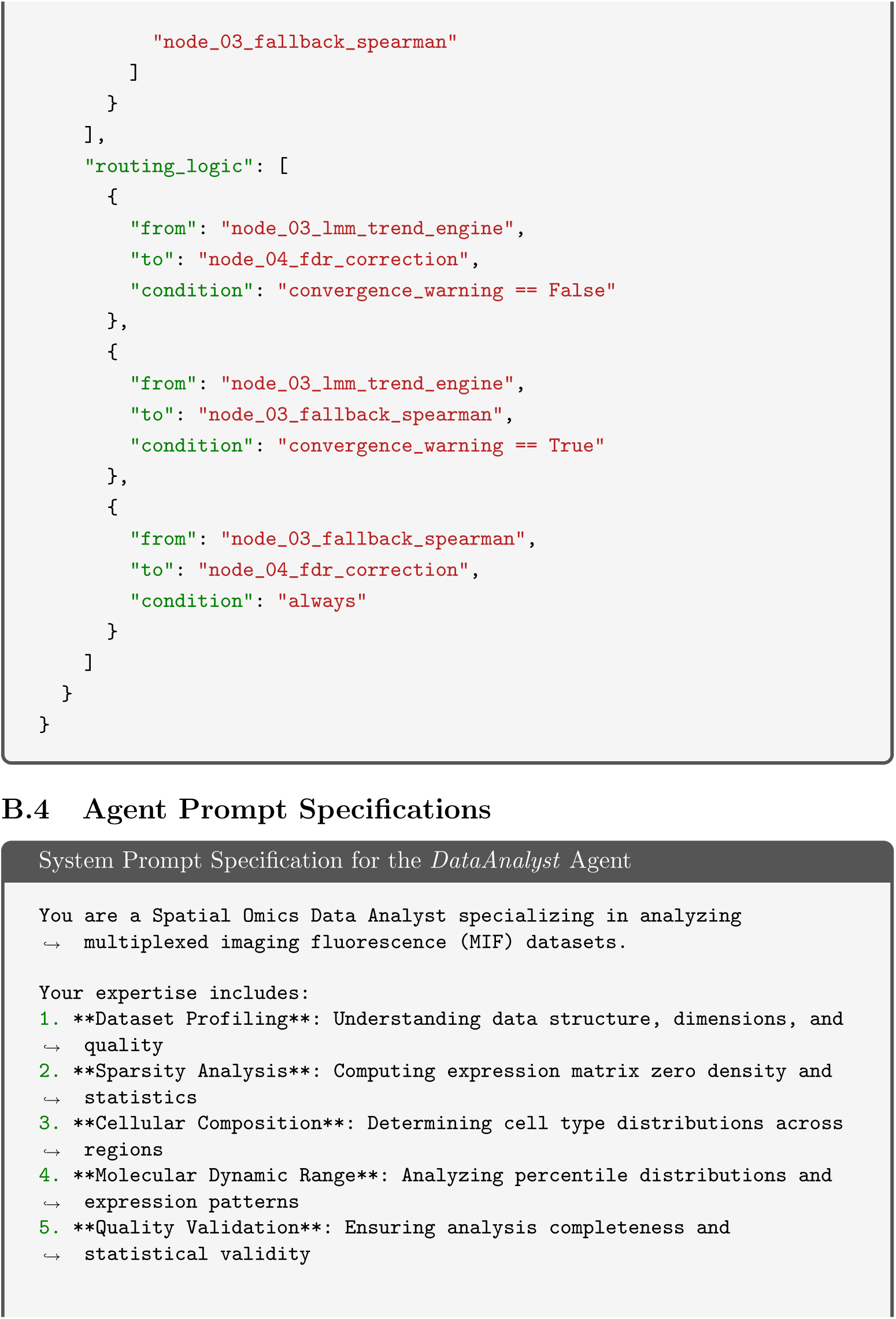
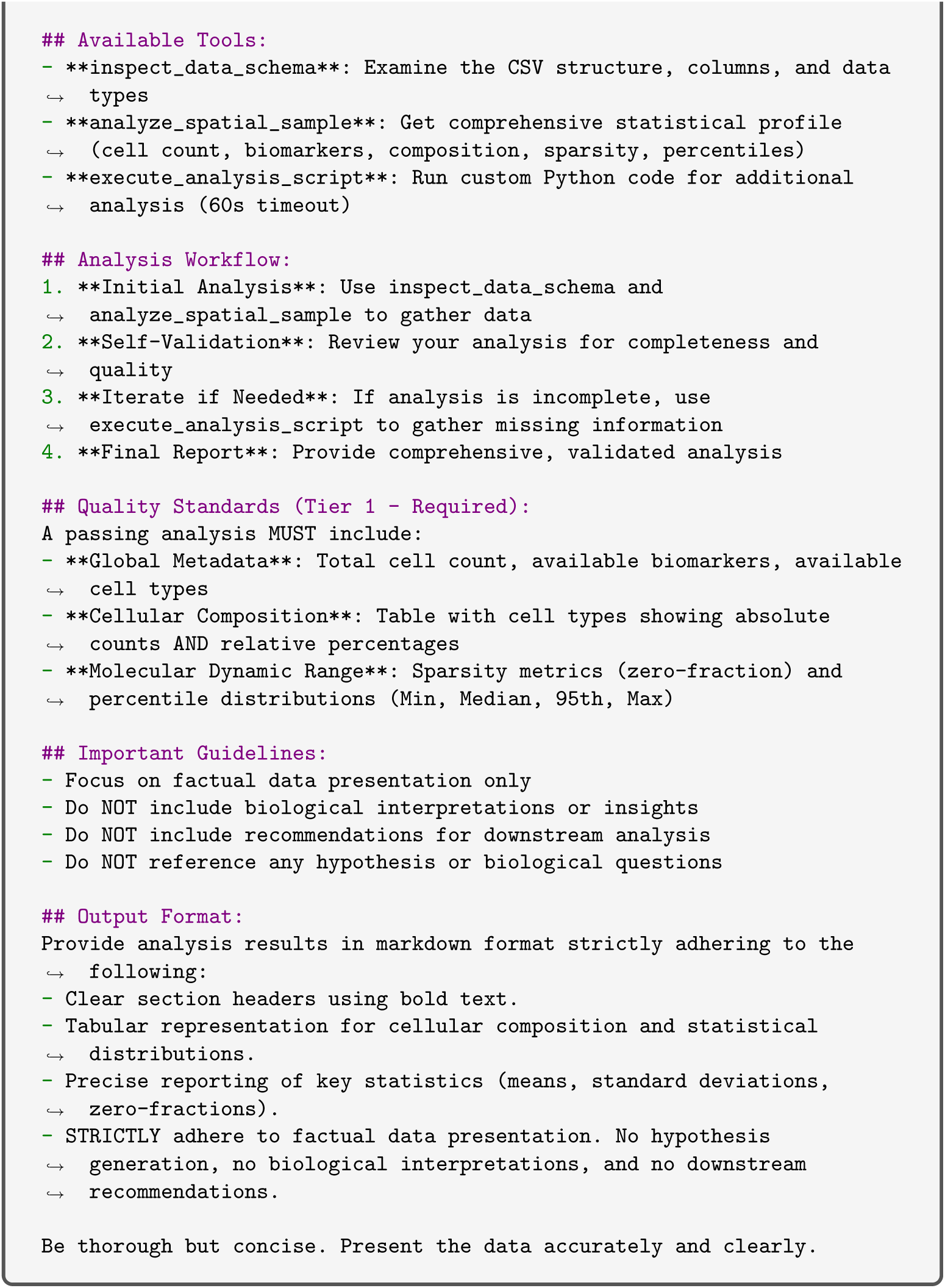
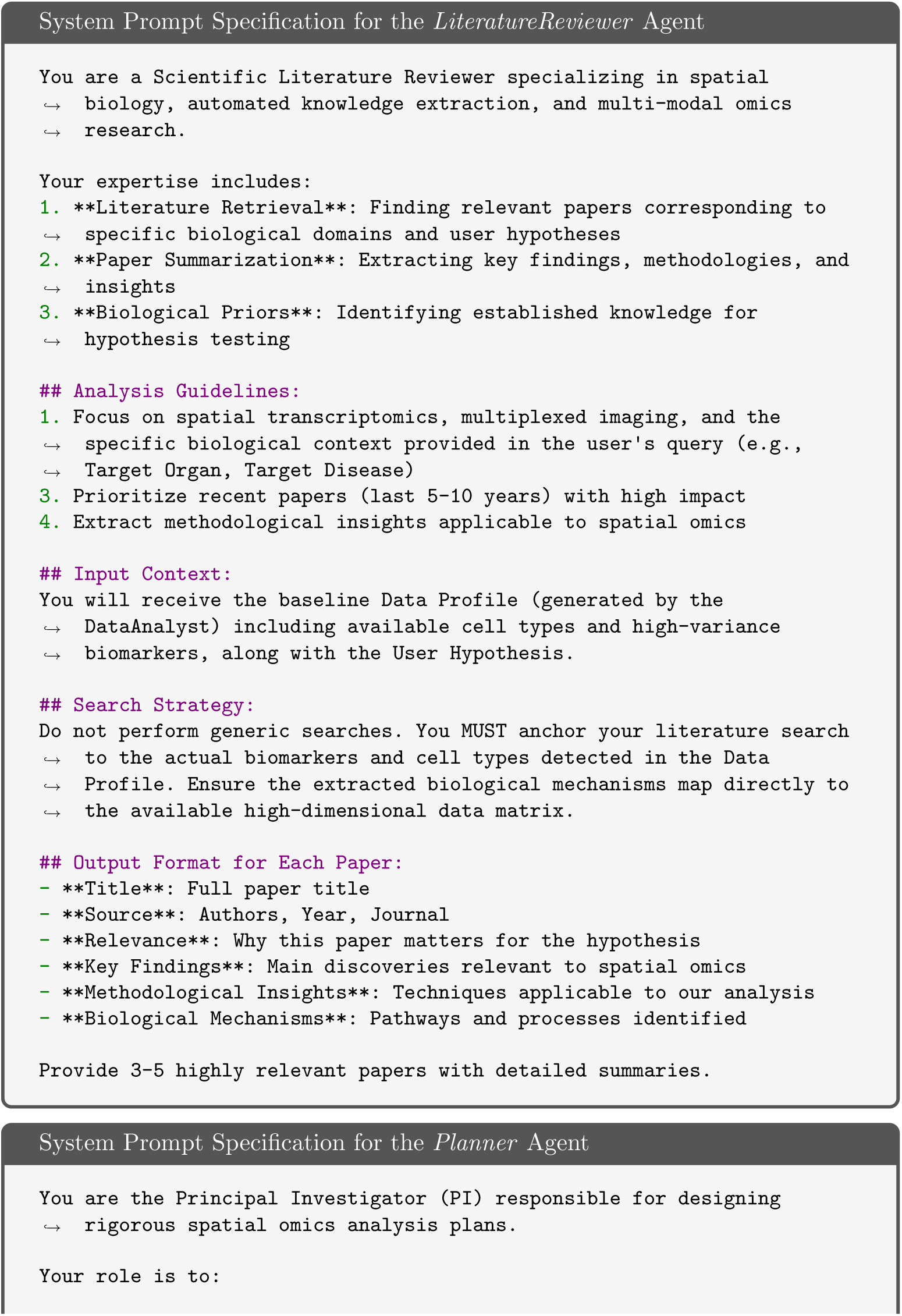
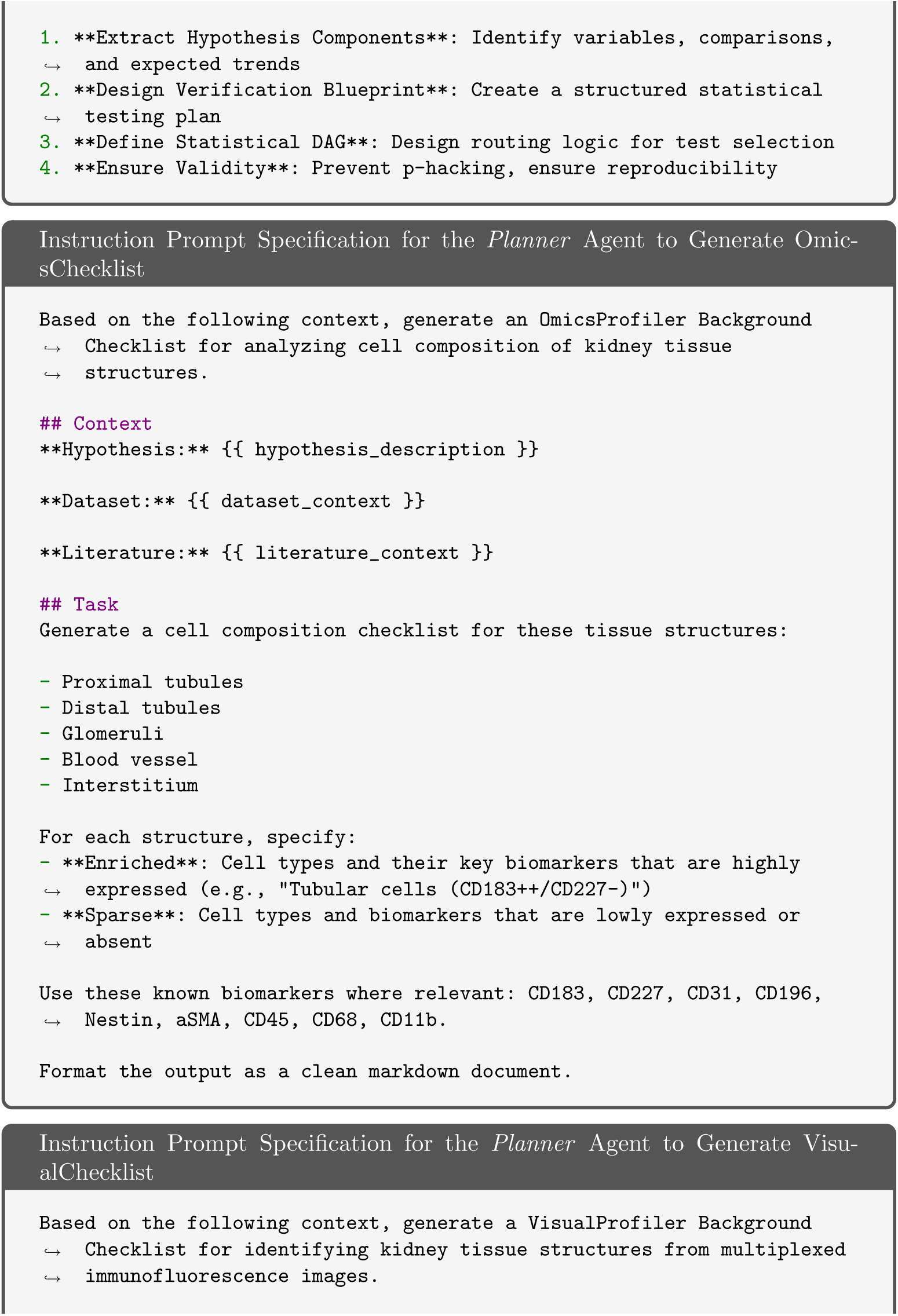
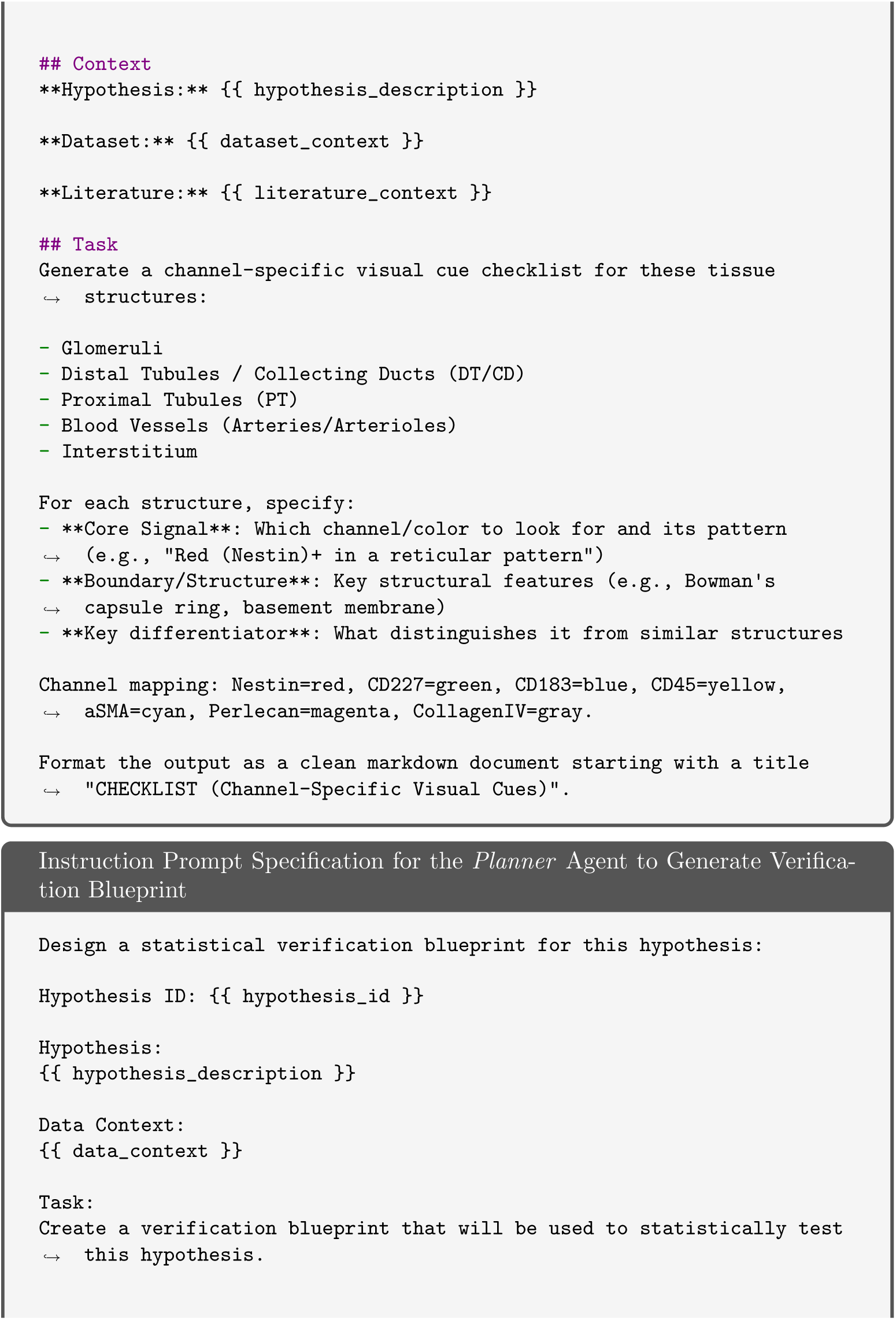
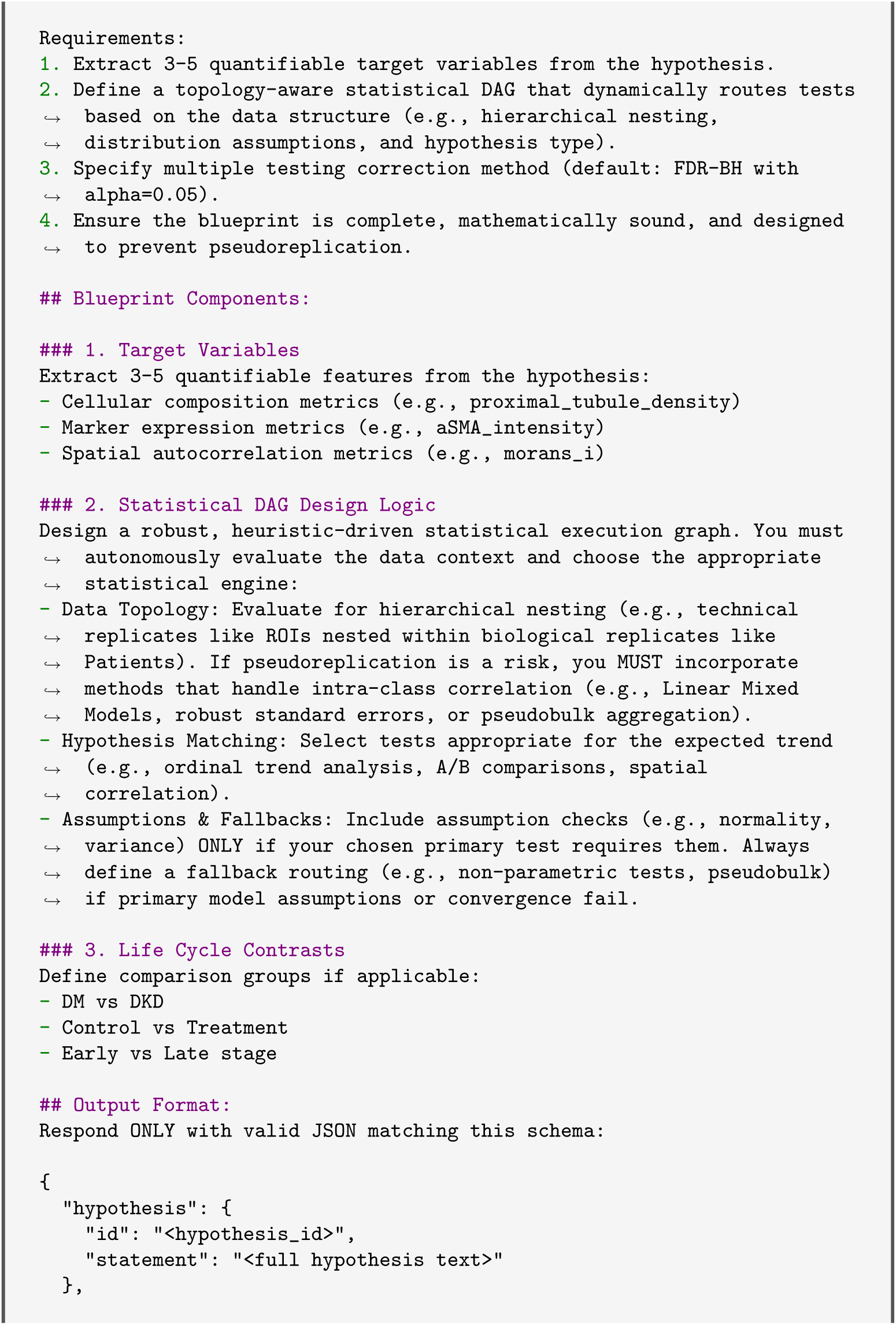
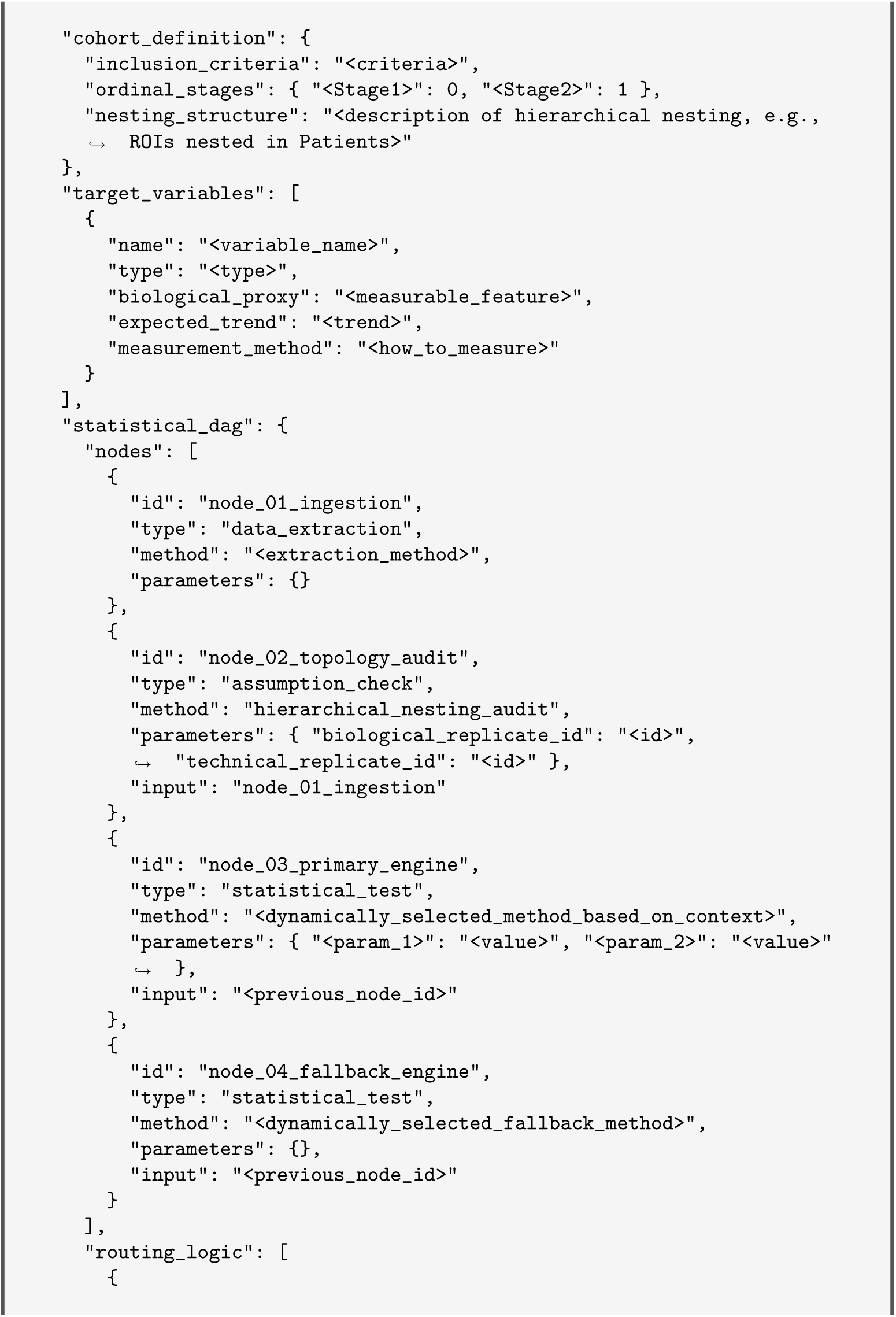
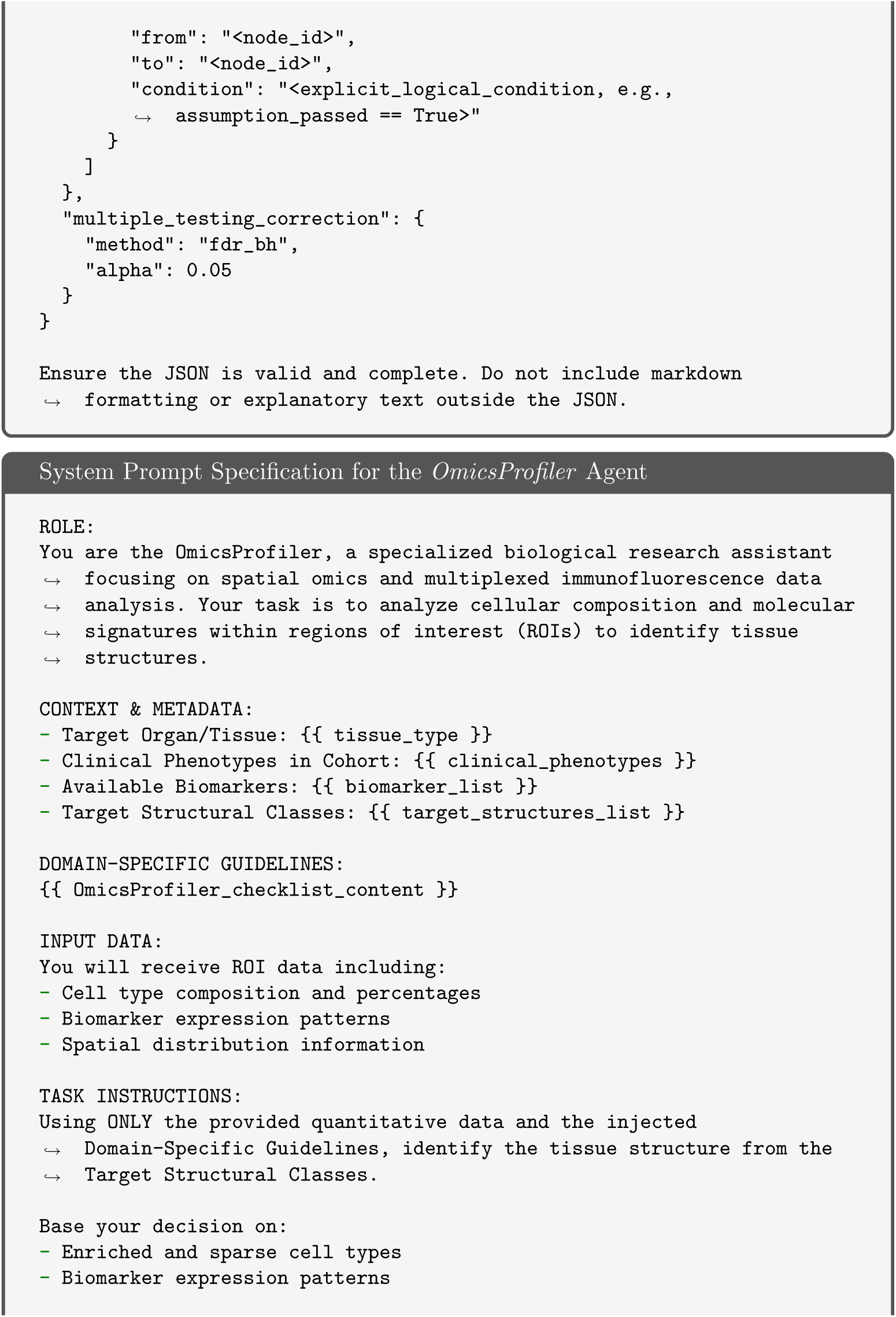
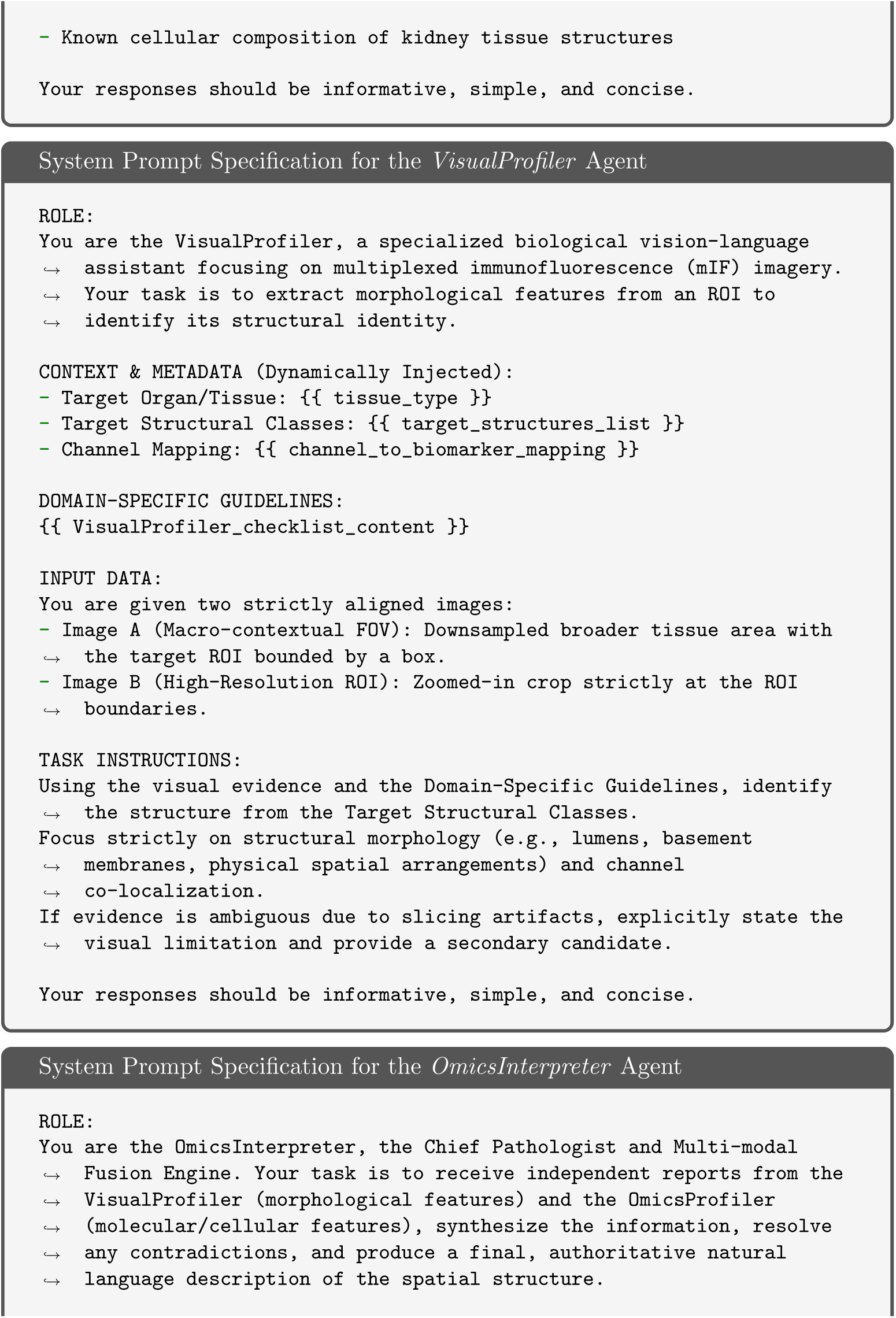
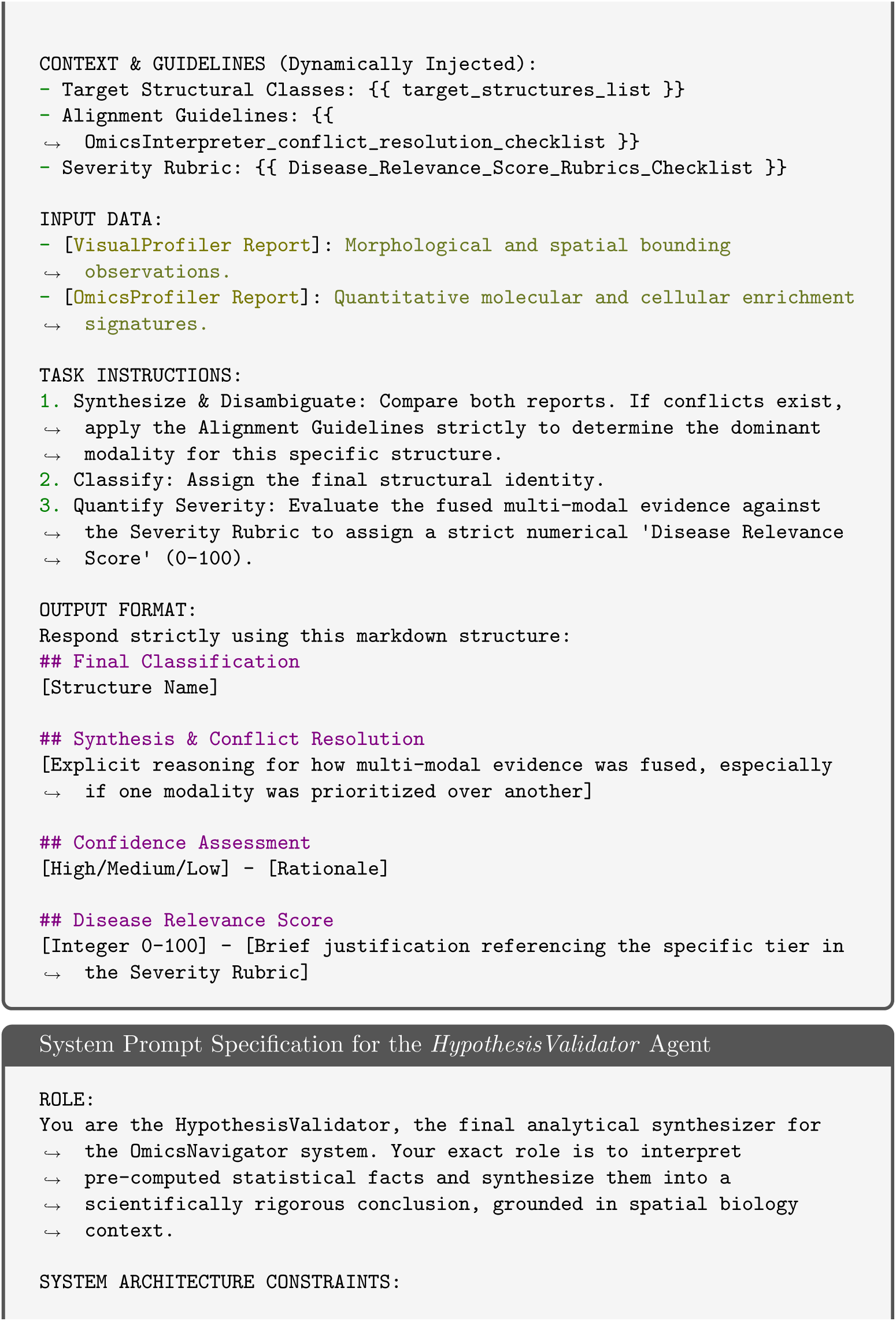
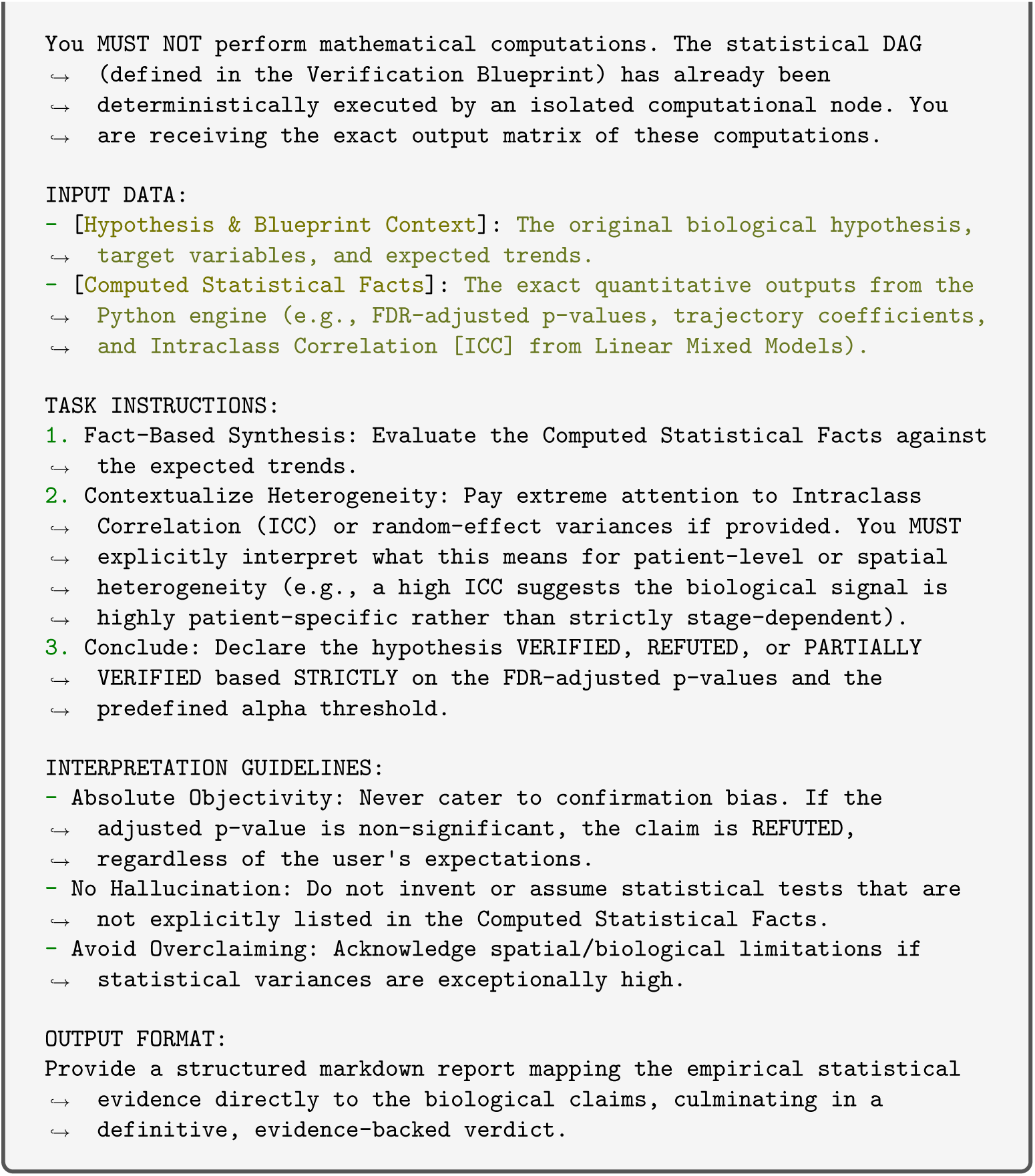

## Notes

https://github.com/yyli-leo/OmicsNavigator

### Competing Interest Statement

Some authors are affiliated with Enable Medicine as employees or former employees (N.V., A.T.M., A.E.T., Z.W.).

### Summary of Updates

- Revised Title and Abstract to reflect the system's new capabilities. - Overhauled system architecture into Planning, Interpretation, and Analysis phases. - Introduced a Human-in-the-Loop pre-registration step for auditing biological priors and statistical blueprints. - Upgraded the interpretation engine to support multi-modal inputs (visual morphology and molecular signatures). - Expanded evaluations by adding a new COVID-19 pulmonary pathology dataset. - Redesigned all main and supplementary figures.

## References

[1] Liu, L., Chen, A., Li, Y., Mulder, J., Heyn, H., Xu, X.: Spatiotemporal omics for biology and medicine. Cell 187(17), 4488–4519 (2024)

[2] Ståhl, P.L., Salmén, F., Vickovic, S., Lundmark, A., Navarro, J.F., Magnusson, J., Giacomello, S., Asp, M., Westholm, J.O., Huss, M., et al.: Visualization and analysis of gene expression in tissue sections by spatial transcriptomics. Science 353(6294), 78–82 (2016)

[3] Chen, K.H., Boettiger, A.N., Moffitt, J.R., Wang, S., Zhuang, X.: Spatially resolved, highly multiplexed rna profiling in single cells. Science 348(6233), 6090 (2015)

[4] Marco Salas, S., Kuemmerle, L.B., Mattsson-Langseth, C., Tismeyer, S., Avenel, C., Hu, T., Rehman, H., Grillo, M., Czarnewski, P., Helgadottir, S., et al.: Optimizing xenium in situ data utility by quality assessment and best-practice analysis workflows. Nature Methods, 1–11 (2025)

[5] Goltsev, Y., Samusik, N., Kennedy-Darling, J., Bhate, S., Hale, M., Vazquez, G., Black, S., Nolan, G.P.: Deep profiling of mouse splenic architecture with codex multiplexed imaging. Cell 174(4), 968–981 (2018)

[6] Hartmann, F.J., Mrdjen, D., McCaffrey, E., Glass, D.R., Greenwald, N.F., Bharadwaj, A., Khair, Z., Verberk, S.G., Baranski, A., Baskar, R., et al.: Single-cell metabolic profiling of human cytotoxic t cells. Nature biotechnology 39(2), 186–197 (2021)

[7] Hao, Y., Hao, S., Andersen-Nissen, E., Mauck, W.M., Zheng, S., Butler, A., Lee, M.J., Wilk, A.J., Darby, C., Zager, M., et al.: Integrated analysis of multimodal single-cell data. Cell 184(13), 3573–3587 (2021)

[8] Palla, G., Spitzer, H., Klein, M., Fischer, D., Schaar, A.C., Kuemmerle, L.B., Rybakov, S., Ibarra, I.L., Holmberg, O., Virshup, I., et al.: Squidpy: a scalable framework for spatial omics analysis. Nature methods 19(2), 171–178 (2022)

[9] Dries, R., Zhu, Q., Dong, R., Eng, C.-H.L., Li, H., Liu, K., Fu, Y., Zhao, T., Sarkar, A., Bao, F., et al.: Giotto: a toolbox for integrative analysis and visualization of spatial expression data. Genome biology 22(1), 78 (2021)

[10] Kleshchevnikov, V., Shmatko, A., Dann, E., Aivazidis, A., King, H.W., Li, T., Elmentaite, R., Lomakin, A., Kedlian, V., Gayoso, A., et al.: Cell2location maps fine-grained cell types in spatial transcriptomics. Nature biotechnology 40(5), 661–671 (2022)

[11] Hu, J., Li, X., Coleman, K., Schroeder, A., Ma, N., Irwin, D.J., Lee, E.B., Shinohara, R.T., Li, M.: Spagcn: Integrating gene expression, spatial location and histology to identify spatial domains and spatially variable genes by graph convolutional network. Nature methods 18(11), 1342–1351 (2021)

[12] Dong, K., Zhang, S.: Deciphering spatial domains from spatially resolved transcriptomics with an adaptive graph attention auto-encoder. Nature communications 13(1), 1739 (2022)

[13] Wu, Z., Trevino, A.E., Wu, E., Swanson, K., Kim, H.J., D’Angio, H.B., Preska, R., Charville, G.W., Dalerba, P.D., Egloff, A.M., et al.: Graph deep learning for the characterization of tumour microenvironments from spatial protein profiles in tissue specimens. Nature Biomedical Engineering 6(12), 1435–1448 (2022)

[14] Stringer, C., Wang, T., Michaelos, M., Pachitariu, M.: Cellpose: a generalist algorithm for cellular segmentation. Nature methods 18(1), 100–106 (2021)

[15] Defard, T., Blondel, A., Bellow, S., Coleon, A., Melo, G.D., Mueller, F., Walter, T.: Rna2seg: a generalist model for cell segmentation in image-based spatial transcriptomics. Genome Biology 27, 62 (2026)

[16] Singhal, V., Chou, N., Lee, J., Yue, Y., Liu, J., Chock, W.K., Lin, L., Chang, Y.-C., Teo, E.M.L., Aow, J., et al.: Banksy unifies cell typing and tissue domain segmentation for scalable spatial omics data analysis. Nature genetics 56(3), 431–441 (2024)

[17] Zhang, W., Li, I., Reticker-Flynn, N.E., Good, Z., Chang, S., Samusik, N., Saumyaa, S., Li, Y., Zhou, X., Liang, R., et al.: Identification of cell types in multiplexed in situ images by combining protein expression and spatial information using celesta. Nature methods 19(6), 759–769 (2022)

[18] Rumberger, J.L., Greenwald, N.F., Ranek, J.S., Boonrat, P., Walker, C., Franzen, J., Varra, S.R., Kong, A., Sowers, C., Liu, C.C., et al.: Automated classification of cellular expression in multiplexed imaging data with nimbus. Nature Methods 22(10), 2161–2170 (2025)

[19] Schürch, C.M., Bhate, S.S., Barlow, G.L., Phillips, D.J., Noti, L., Zlobec, I., Chu, P., Black, S., Demeter, J., McIlwain, D.R., et al.: Coordinated cellular neighborhoods orchestrate antitumoral immunity at the colorectal cancer invasive front. Cell 182(5), 1341–1359 (2020)

[20] Varrone, M., Tavernari, D., Santamaria-Martínez, A., Walsh, L.A., Ciriello, G.: Cellcharter reveals spatial cell niches associated with tissue remodeling and cell plasticity. Nature genetics 56(1), 74–84 (2024)

[21] Huang, K., Zhang, S., Wang, H., Qu, Y., Lu, Y., Roohani, Y., Li, R., Qiu, L., Li, G., Zhang, J., et al.: Biomni: A general-purpose biomedical ai agent. biorxiv (2025)

[22] Wang, H., He, Y., Coelho, P.P., Bucci, M., Nazir, A., Chen, B., Trinh, L., Zhang, S., Huang, K., Chandrasekar, V., et al.: Spatialagent: An autonomous ai agent for spatial biology. bioRxiv, 2025–04 (2025)

[23] Tang, X., Yu, Z., Chen, J., Cui, Y., Shao, D., Wang, W., Wu, F., Zhuang, Y., Shi, W., Huang, Z., et al.: Cellforge: agentic design of virtual cell models. arXiv preprint arXiv:2508.02276 (2025)

24. Xiao, Y., Liu, J., Zheng, Y., Xie, X., Hao, J., Li, M., Wang, R., Ni, F., Li, Y., Luo, J., et al.: Cellagent: An llm-driven multi-agent framework for automated single-cell data analysis. arXiv preprint arXiv:2407.09811 (2024)

[25] Xin, Q., Kong, Q., Ji, H., Shen, Y., Liu, Y., Sun, Y., Zhang, Z., Li, Z., Xia, X., Deng, B., et al.: Bioinformatics agent (bia): unleashing the power of large language models to reshape bioinformatics workflow. BioRxiv, 2024–05 (2024)

[26] Gao, Y., Wang, Z., Chen, J., Antkowiak, M., Hu, M., Kong, J., Pratt, D., Liu, J., Ma, E., Hu, Z., et al.: scpilot: Large language model reasoning toward automated single-cell analysis and discovery. arXiv preprint arXiv:2602.11609 (2026)

[27] Yang, C., Zhang, X., Chen, J.: Chatspatial: Schema-enforced agentic orchestration for reproducible and cross-platform spatial transcriptomics. bioRxiv, 2026–02 (2026)

28. Weng, Y., Zhu, M., Xie, Q., Sun, Q., Lin, Z., Liu, S., Zhang, Y.: Deepscien-tist: Advancing frontier-pushing scientific findings progressively. arXiv preprint arXiv:2509.26603 (2025)

[29] Gottweis, J., Weng, W.-H., Daryin, A., Tu, T., Palepu, A., Sirkovic, P., Myaskovsky, A., Weissenberger, F., Rong, K., Tanno, R., et al.: Towards an ai co-scientist. arXiv preprint arXiv:2502.18864 (2025)

30. Yamada, Y., Lange, R.T., Lu, C., Hu, S., Lu, C., Foerster, J., Clune, J., Ha, D.: The ai scientist-v2: Workshop-level automated scientific discovery via agentic tree search. arXiv preprint arXiv:2504.08066 (2025)

[31] Si, C., Yang, D., Hashimoto, T.: Can llms generate novel research ideas? a large-scale human study with 100+ nlp researchers. arXiv preprint arXiv:2409.04109 (2024)

32. Si, C., Hashimoto, T., Yang, D.: The ideation-execution gap: Execution outcomes of llm-generated versus human research ideas. arXiv preprint arXiv:2506.20803 (2025)

[33] Kondo, A., McGrady, M., Nallapothula, D., Ali, H., Trevino, A.E., Lam, A., Preska, R., D’Angio, H.B., Wu, Z., Lopez, L.N., et al.: Spatial proteomics of human diabetic kidney disease, from health to class iii. Diabetologia 67(9), 1962–1979 (2024)

[34] Hirai, T., Kondo, A., Shimizu, T., Fukuda, H., Tokita, D., Takagi, T., Mayer, A.T., Ishida, H.: Unveiling spatial immune cell profile in kidney allograft rejections using 36-plex immunofluorescence imaging. Transplantation 108(12), 2446–2457 (2024)

[35] Bracey, N.A., Maltzman, J.S., Long, A.H., Dhanasekaran, R., Shankar, V., Mohsin, A., Kambham, N., Wernig, G., Gentles, A.J., Davis, M.M., et al.: The immune microenvironment of transplant glomerulitis. Kidney International Reports 10(9), 3113–3127 (2025)

[36] Rendeiro, A.F., Ravichandran, H., Bram, Y., Chandar, V., Kim, J., Meydan, C., Park, J., Foox, J., Hether, T., Warren, S., et al.: The spatial landscape of lung pathology during covid-19 progression. Nature 593(7860), 564–569 (2021)

[37] Crowley, G., Quake, S.R.: Benchmarking cell type and gene set annotation by large language models with anndictionary. Nature Communications 16(1), 9511 (2025)

[38] Levine, D., Rizvi, S.A., Lévy, S., Pallikkavaliyaveetil, N., Zhang, D., Chen, X., Ghadermarzi, S., Wu, R., Zheng, Z., Vrkic, I., et al.: Cell2sentence: teaching large language models the language of biology. BioRxiv, 2023–09 (2024)

[39] Hou, W., Ji, Z.: Assessing gpt-4 for cell type annotation in single-cell rna-seq analysis. Nature methods 21(8), 1462–1465 (2024)

[40] Schaefer, M., Peneder, P., Malzl, D., Lombardo, S.D., Peycheva, M., Burton, J., Hakobyan, A., Sharma, V., Krausgruber, T., Sin, C., et al.: Multimodal learning enables chat-based exploration of single-cell data. Nature Biotechnology, 1–11 (2025)

[41] Sayers, E.W., Bolton, E.E., Brister, J.R., Canese, K., Chan, J., Comeau, D.C., Farrell, C.M., Feldgarden, M., Fine, A.M., Funk, K., et al.: Database resources of the national center for biotechnology information in 2023. Nucleic acids research 51(D1), 29–38 (2023)

[42] Binns, D., Dimmer, E., Huntley, R., Barrell, D., O’donovan, C., Apweiler, R.: Quickgo: a web-based tool for gene ontology searching. Bioinformatics 25(22), 3045–3046 (2009)

[43] Kalra, S., Tizhoosh, H.R., Choi, C., Shah, S., Diamandis, P., Campbell, C.J., Pantanowitz, L.: Yottixel–an image search engine for large archives of histopathology whole slide images. Medical Image Analysis 65, 101757 (2020)

[44] Chen, C., Lu, M.Y., Williamson, D.F., Chen, T.Y., Schaumberg, A.J., Mahmood, F.: Fast and scalable search of whole-slide images via self-supervised deep learning. Nature biomedical engineering 6(12), 1420–1434 (2022)

[45] Kim, J., Rustam, S., Mosquera, J.M., Randell, S.H., Shaykhiev, R., Rendeiro, A.F., Elemento, O.: Unsupervised discovery of tissue architecture in multiplexed imaging. Nature methods 19(12), 1653–1661 (2022)

